# Operative dimensions in unconstrained connectivity of recurrent neural networks

**DOI:** 10.1101/2022.06.03.494670

**Authors:** Renate Krause, Matthew Cook, Sepp Kollmorgen, Valerio Mante, Giacomo Indiveri

**Author notes:** Equal contribution (Valerio Mante, Giacomo Indiveri).

## Abstract

Recurrent Neural Networks (RNNs) are commonly used models to study neural computation. However, a comprehensive understanding of how dynamics in RNNs emerge from the underlying connectivity is largely lacking. Previous work derived such an understanding for RNNs fulfilling very specific constraints on their connectivity, but it is unclear whether the resulting insights apply more generally. Here we study how network dynamics are related to network connectivity in RNNs trained without any specific constraints on several tasks previously employed in neuroscience. Despite the apparent high-dimensional connectivity of these RNNs, we show that a low-dimensional, functionally relevant subspace of the weight matrix can be found through the identification of *operative* dimensions, which we define as components of the connectivity whose removal has a large influence on local RNN dynamics. We find that a weight matrix built from only a few operative dimensions is sufficient for the RNNs to operate with the original performance, implying that much of the high-dimensional structure of the trained connectivity is functionally irrelevant. The existence of a low-dimensional, operative subspace in the weight matrix simplifies the challenge of linking connectivity to network dynamics and suggests that independent network functions may be placed in specific, separate subspaces of the weight matrix to avoid catastrophic forgetting in continual learning.

## 1 Introduction

A central goal in neuroscience is to understand how groups of tightly interconnected neurons generate the complex network dynamics that underlies behavior. To this end, ever larger experimental datasets on neural anatomy, neural activity and the corresponding behavior are collected and analyzed and new experimental tools to extend the amount and quality of such data are continuously developed. However, in most settings it remains an open question how the underlying neural connectivity is able to generate the observed neural dynamics. Progress in how to inspect and interpret these complex datasets, and in particular on the relation between neural structure and function, may come in particular from new theoretical frameworks [1, 2].

Artificial Recurrent Neural Networks (RNNs) are a promising tool to develop such theoretical frameworks in a well-controlled and flexible setting [3, 4, 5]. Previous work on RNNs explained how a specifically designed connectivity can give rise to desired network dynamics [6, 7, 8]. Further theoretical work on RNNs with random, recurrent weights provided detailed insights into the properties of neural dynamics emerging from largely unstructured connectivity [9, 10, 11, 12, 13]. More recent work related structure to function in feedforward networks [14, 15] or RNNs with specific connectivity constraints. For example, network motifs in threshold-linear networks are used to predict the existence of fixed points of the dynamics [16]. Similarly, a principled understanding of dynamics and the role of different cell classes in computations can be achieved for RNNs with low-dimensional weight matrices (low-rank RNN [17, 18]). At present, it remains unclear if and how the resulting findings can be generalized to RNNs which are not subject to such constraints.

In this work, we study how the network dynamics are related to the network connectivity in vanilla RNNs that are trained using a gradient-based approach without imposing any specific constraints on the network weight matrix. We find that the weight matrix of the trained RNN is consistently highdimensional, even when trained on tasks resulting in dynamics that are low-dimensional. Notably, we are nonetheless able to identify a low-dimensional subspace within the high-dimensional weight matrix that is sufficient to perform the trained task. We identify this functionally relevant subspace of the connectivity through the definition of a set of *operative dimensions*, which we define as components of the network connectivity that have a large impact on computationally relevant local dynamics produced by the network.

This ability to identify functionally relevant subspaces in weight matrices improves our understanding of how the network connectivity generates the observed network dynamics and thereby makes RNNs into a more interpretable model for neuroscience and machine learning applications.

## 2 Results

We perform our analyses on vanilla RNNs trained without regularization terms, using the standard RNN equation:

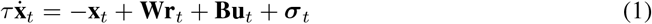

where 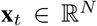 are the linear activities of the *N* hidden units over time *t* with 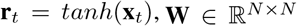 is the recurrent weight matrix of the hidden units and 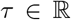 is the time constant (*τ* = 10*ms, dt* = 1*ms*). We consider RNNs of *N* = 100 noisy units, where each element of ***σ**_t_* is drawn from a Gaussian distribution 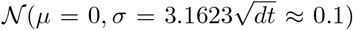. The network output is defined as:

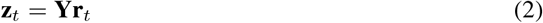

with output readout weights 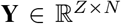. The task-dependent, time-varying inputs 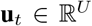 are projected onto the hidden units with input weights 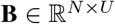. Note that only the inputs vary across different conditions within a task. For any given condition, the cost is defined as:

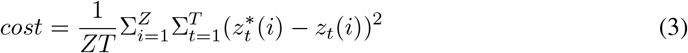

where 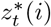 is the desired output. All network weights (**B**, **W**, **Y**) are randomly initialized, and networks are trained to minimize the summed costs across all conditions.

The RNNs are trained separately on two previously proposed tasks: context-dependent integration [19] and sine wave generation [20] (see appendix section A.3.8 for additional results on sequential MNIST). The dynamics of RNNs trained on these tasks is well understood, and was shown to be largely independent of the type of employed RNN [21]. In context-dependent integration, the RNN receives two noisy, sensory inputs (between −1 and 1) and two static, context inputs (0 or 1). The network is trained to select one of the two sensory inputs (depending on the currently active context input; i.e. select input sensory_i_ in context_i_) and integrate it over time (Fig. 1a, b). The network should reach choice_1_ or choice_2_ if the average of the contextually relevant sensory input is positive or negative, respectively. In the sine wave generation task, the RNN receives one static input and is trained to output a sine wave whose target frequency is given by the level of this static input (Fig. 1f, g; for details on task structures see section A.1.1). For both tasks we trained 20 RNNs (different random initial connectivity) with gradient-based optimization to minimize the cost (Eq. 3; for details see section A.1.2).

**Figure 1:**
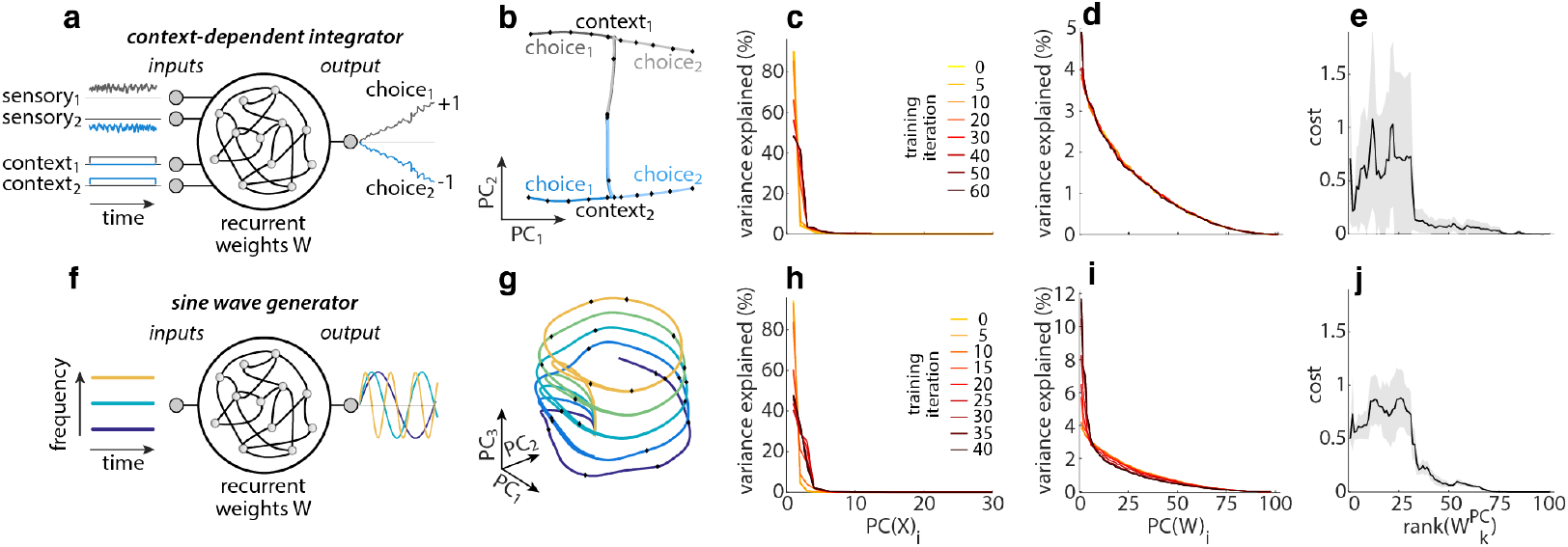
High-variance dimensions of the connectivity. (**a**) Task schematic of the context-dependent integrator. (**b**) Low-dimensional projection of condition average trajectories for an example contextdependent integrator. (**c**) Variance explained (in activity space) by individual PCs of the network activity **X** over all input conditions, shown at different stages of training. (**d**) Variance explained (in weight space) by individual PCs of the weight matrix **W** at different stages of training. (**e**) Network output cost (Eq. 3) of networks with reduced-rank weight matrices 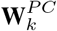 for *k* = 1 : *N* (Eq. 5), averaged over input conditions (shaded area: median absolute deviation (*mad*) over trials). (**f-j**) Analogous to (**a-e**), but for a sine wave generator. (**b-e**) and (**g-j**) 1 representative network per task.

### 2.1 High-variance dimensions

To characterize the relation between network connectivity and dynamics, we first consider the functional relevance of *high-variance* dimensions (as in analyses of low-rank RNN [17]) defined as the left singular vectors of the RNN weight matrix **W**:

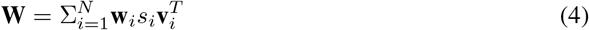

where **w**_*i*_ and **v**_*i*_ are the left and right singular vectors respectively, and *s_i_* the associated singular values. In general, we refer to the i-th left singular vectors of a matrix 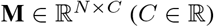 as the principal components of the matrix, *PC*(**M**)_*i*_.

We first ask whether the weight matrices of the trained networks are low or high-dimensional, by assessing the amount of variance in the connectivity 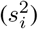 explained by principal components *PC*(**W**)_*i*_. In the trained networks, variance explained falls off slowly over *PC*(**W**)_*i*_, implying that the weight matrices **W** are high-dimensional. The variance explained by individual PCs changes little during training (Fig. 1d, *i* shown for column dimensions; colors: training iteration) but is strongly influenced by the rank of the untrained network (Fig. S6). Despite having a high-dimensional connectivity, the network activities (**X** = [**x**_1_, … **x**_*T*_] and **R** = [**r**_1_, … **r**_*T*_]; both further concatenated over all input conditions) are consistently low-dimensional. Activity is low-dimensional even before training, as **W** is initialized with a spectral radius of 1 (Fig. 1c, h; colors: training iteration). Throughout training, more than 95% of the variance in the network activities **X** can be described by less than ten dimensions. Thus, even though the underlying weight matrix (being high-dimensional) could support activity in any region of state space, the network activity generally only occupies a low-dimensional subspace. This finding raises the question of whether all dimensions in **W** are truly necessary to perform the task.

To assess the functional importance of single dimensions of **W**, we construct reduced-rank approximations of **W** from only a subset of all *PC*(**W**)_*i*_ and then assess the performance of the resulting RNN (as in [22]). Each low-rank approximation is constructed by including only the first *k PC*(**W**)_*i*_:

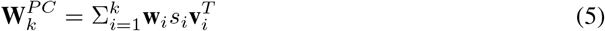

RNN performance based on 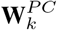 is evaluated as above to obtain the network output cost (Eq. 3).

We find that a large number of *PC*(**W**)_*i*_ are consistently required to reach a similar performance as the original network. For the two example networks, 88 PCs are required to achieve original performance for context-dependent integration, and 86 PCs for sine wave generation (Fig. 1e, j; we define *original performance* as 4 times the cost of the corresponding full-rank RNN, Fig. S7 shows more example networks). Furthermore, performance does not improve monotonically with increasing rank, but rather displays sudden jumps at specific ranks (Fig. 1e, j). These jumps suggest that some high-variance dimensions are more relevant than others in driving the network output. These putative differences in functional relevance are not obviously mirrored in the amount of variance in **W** explained by each high-variance dimension (Fig. 1d, i). In other words, the amount of variance in connectivity space explained by a given dimension in **W** does not appear to directly correspond to its functional relevance.

Overall, it appears that the inferred high-variance dimensions may be ill-suited to shed light on the relation between RNN connectivity and dynamics. These findings point to two possible scenarios: the functionally relevant subspace of the weight matrix in these unconstrained RNNs is genuinely highdimensional, precluding simpler descriptions of the most relevant components of the connectivity; or high-variance dimensions *PC*(**W**)_*i*_ may in general not be well-aligned with functionally relevant dimensions of the connectivity.

### 2.2 Operative dimensions

#### 2.2.1 Definition of operative dimensions

To distinguish between these two scenarios, we devised an approach to directly identify functionally relevant dimensions of the weight matrix. We refer to this new type of dimensions as the *operative dimensions* of the connectivity.

The definition of operative dimensions is based on the key insight that sudden jumps in performance (*cost*, Eq. 3) can be caused by small changes in the local dynamics. Here we illustrate this for one example network trained on context-dependent integration (Fig. 2). We focus on 2 out of 8 input conditions, for which the cost shows prominent jumps at specific ranks *k*, specifically when transitioning from *k* = 60 to *k* = 61 (Fig. 2a; red circles).

**Figure 2:**
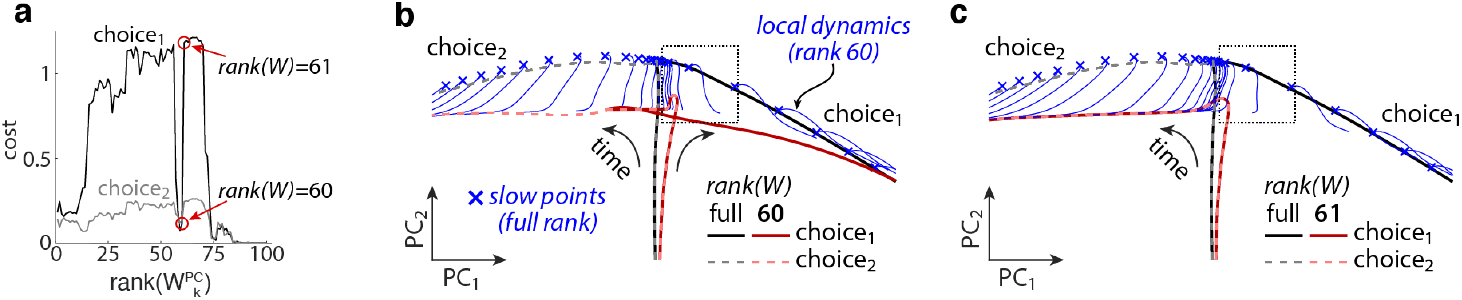
Local and global dynamics. (**a**) Network output cost for reduced-rank weight matrices 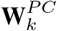 in an example network for context-dependent integration. Analogous to Fig. 1**e**, but shown separately for choice_1_ and choice_2_ trials in context_1_. (**b**) Average trajectories for the conditions in (**a**) in the full-rank and a reduced-rank (*k* = 60) network (grayish and reddish curves, see legend). Blue lines show recurrent dynamics in the reduced-rank network initialized at locations corresponding to slow points of the dynamics in the full-rank network (blue crosses). (**c**) Same as (**b**) but for a reduced-rank network with *k* = 61.

To understand this sharp increase, we examine the activity trajectories produced by the original full-rank and the reduced-rank RNN for the two considered input conditions (*k* = 60 in Fig. 2b; *k* = 61 in Fig. 2c). The well-performing, full-rank RNN produces trajectories that start in the center and move out to endpoints on the right (*choice_1_*; solid black) or left (*choice_2_*; dashed gray). The RNN with *k* = 60 closely matches these dynamics (Fig. 2b; red curves, solid and dashed). On the other hand, the RNN with *k* = 61 erroneously produces trajectories that move to the left endpoint for both conditions (Fig. 2c; overlapping red curves). This discrepancy between *choice_1_* trajectories in the original full-rank and *k* = 61 RNN underlies the large jump in cost (Fig. 2a).

Critically, we found that the large difference in cost between the two reduced-rank networks is due to only a small change in the underlying local, recurrent dynamics. To visualize the local dynamics in the two reduced-rank RNNs, we generate brief activity trajectories (Fig. 2b,c; blue curves) initialized at the location of identified slow-points of the dynamics in the original full-rank RNN (blue crosses; identified as in [19]). Here, we isolate the recurrent contribution to the dynamics by running the RNN without any external inputs (Eq. 1 with **u**_*t*_ = 0, ***σ**_t_* = 0). The resulting trajectories suggest that the local recurrent dynamics is substantially different across networks only in the region of state space that activity travels through at the onset of the trial (Fig. 2b,c; dashed square). This small local difference is sufficient to redirect the *choice_1_* trajectories for *k* = 61 towards the wrong endpoint. This mistake cannot be corrected by recurrent dynamics at other locations in state space, resulting in the large cost. In summary, the functional relevance of individual dimensions of **W** is hard to assess based on changes in the cost, but rather may be better judged based on its effect on local network dynamics.

Based on this insight, we define *operative dimensions* as dimensions in **W** that have a large impact on the local dynamics if removed from **W** (Fig. 3a). Given an arbitrary unit vector 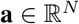, we define 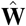 as the matrix of rank *N* – 1 obtained by removing the dimension **a** from the column space of **W** (orthogonal projection):

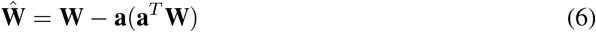

**Figure 3:**
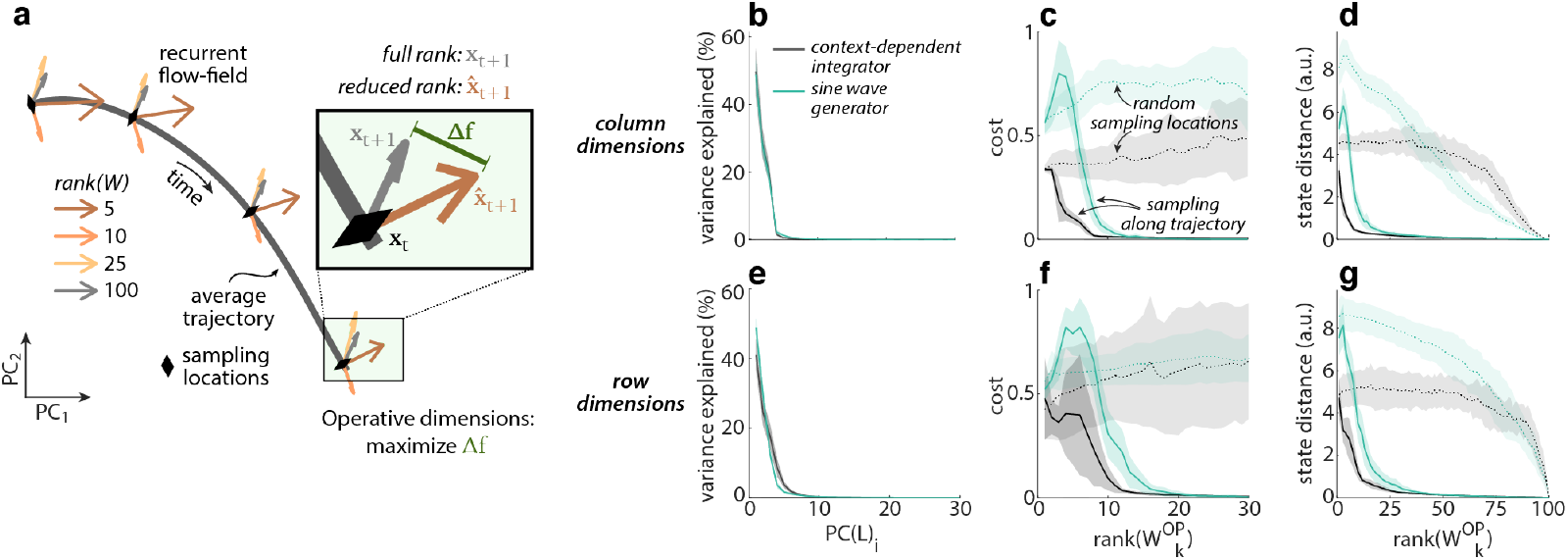
Operative dimensions. (**a**) Definition of operative dimensions based on local recurrent dynamics along a condition average trajectory. Arrows show the recurrent contribution to the dynamics for the full-rank and several reduced-rank networks (colors, see legend). Local operative dimensions maximize Δ*f*. Here inputs **u**_*t*_ and noise **σ**_*t*_ are set to zero. (**b**) Rank of global operative column dimensions, estimated with PC analysis on concatenated local operative dimensions (Eq. 10 and 11). (**c**) Network output cost of networks with reduced-rank weight matrix 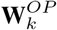 for *k* = 1 : *N* (Eq. 12). (**d**) State distance between trajectories in the full-rank network and in networks with reduced-rank weight matrix 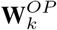. (**b-d**) Based on global operative *column* dimensions and averaged over 20 networks per task; shaded area: *mad*. Network output cost obtained with internal and input noise, state distance without any noise. (**e-g**) Same as (**b-d**) for global operative *row* dimensions.

In the absence of noise, the dynamics of the resulting reduced-rank RNN are then given by 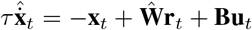. At location **x**_*t*_ the activity evolves to 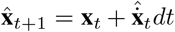 over one time-step. Likewise, the state of the full-rank network evolves to 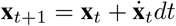, where we have set ***σ**_t_* = 0 in Eq. 1. We quantify the change in local dynamics brought about by removing **a** as:

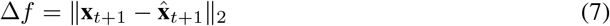

as illustrated graphically in Fig. 3a. We then define the operative dimensions of **W** based on Δ*f* in two steps. In the first step, we infer a set of *local* operative dimensions at each of *P* sampling locations 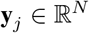 in state space. The sampling locations are chosen as an evenly distributed subset of the states **x**_*t*_ explored by the condition average trajectories of the full-rank network (Fig. 1b, g; only a subset of *P* > 100 locations shown). This choice of sampling locations ensures that we consider only local dynamics that is likely to contribute to solving the task at hand. The first local operative dimension **d**_1, *j*_ at location **y**_*j*_ is defined as the dimension **a** that maximizes Δ*f*:

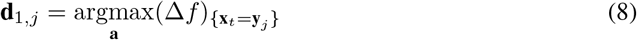

Up to *N* – 1 further local operative dimensions **d**_*i,j*_ at the same location **y**_*j*_ are defined in the same way, but under the additional constraint that they need to be orthogonal to all previously identified local operative dimensions at that location:

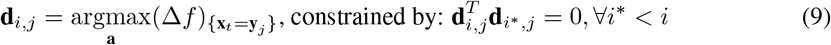

In the second step, we define the *global* operative dimensions by combining the local operative dimensions **d**_*i,j*_ from all sampling locations **y**_*j*_ (*i* = 1 : *N, j* = 1 : *P*). Specifically, the local operative dimensions are scaled by their local Δ*f* and concatenated into one matrix **L**.

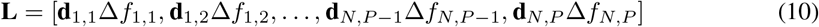

where 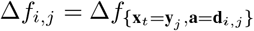. The i-th global operative dimensions **q**_*i*_ is then defined as the i-th left singular vector of **L**:

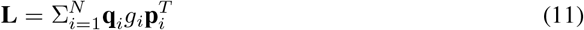

where *g_i_* are the singular values, and **p**_*i*_ the right singular vectors of **L**. The subspace spanned by the global operative dimensions **q**_*i*_ consists of all left singular vectors with *g_i_* > 0. Note that these steps result in operative dimensions of the *column* space of **W**, which are referred to as *column dimensions* in the figures. We employ an analogous approach to define operative dimensions of the *row* space of **W** which yields global operative row dimensions 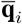 (*row dimensions* in figures; see section A.2.2).

#### 2.2.2 Operative dimensions identify a low-d functional subspace of the connectivity

To quantify the functional relevance of the global operative dimensions, we proceed as for the high-variance dimensions above (Eq. 5) by constructing reduced-rank approximations of **W** from only a subset of the operative dimensions. For the column dimensions, the reduced-rank approximations 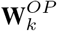 are given by:

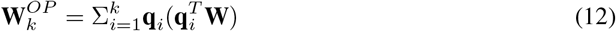

Network activity 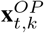 based on 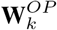 is then computed as above (Eq. 1).

Unlike the high-variance dimensions, the global operative dimensions identify a low-dimensional subspace that is sufficient for the RNN to achieve the original performance in both tasks (Fig. 3b-c, e-f; solid lines, qualitatively similar results on sequential MNIST are shown in section A.3.8). We find that 15 column and 27 row dimensions are sufficient to achieve original performance for contextdependent integration; and 29 column and 41 row dimensions for sine wave generation. Thus, the RNNs are *functionally* low-rank even though the underlying weight matrix is high-dimensional.

This clear difference between the global operative dimensions and the high-variance dimensions is also reflected in their weak alignment to each other (Fig. S8). This finding implies that much of the network connectivity in these trained networks is not required to perform the task.

Notably, the identified functionally relevant subspace of the connectivity is sufficient to generate the original network trajectory, not just the network activities along the output direction. The average distance between the network trajectories of the reduced-rank and the full-rank networks decreases rapidly as the global operative dimensions are sequentially added to the weight matrix (Fig. 3d, g; 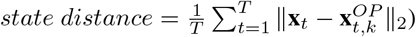. This observation follows from the definition of operative dimensions, which focuses not on changes in network output, but rather on the recurrent dynamics at sampling locations lying along the entirety of the condition average trajectories. We obtained similar results for networks at different stages of training (Fig. S14) and with different types of connectivity and architecture (Fig. S15). Similarly, even though operative dimensions are defined based on how much they alter the local dynamics when removed from **W**, the first few operative dimensions are sufficient to preserve the dominant local, linear dynamics (Fig. S17).

Unlike for high-variance dimensions (Fig. 1e, j), RNN performance and state distance change smoothly with increasing rank of the global operative dimensions (Fig. 3c-d, f-g). However, adding the first few dimensions to the weight matrix often hurts performance. This effect can happen because the global operative dimensions are not sorted based on when they are required during individual trials, but rather by their impact on local network dynamics. Indeed, the first few operative dimensions are mostly required at state-space locations explored late in the trial, but are insufficient to sustain the dynamics required early in the trial (Fig. S21c, f, i, l).

Identifying the operative dimensions critically relies on the correct choice of sampling locations. First, we illustrate the importance of the choice of sampling locations by defining global operative dimensions based on random sampling locations in state space (Fig. 3c, f; dotted lines). For the same choice of rank, the resulting dimensions yield much poorer performance compared to the operative dimensions defined as above, emphasizing the importance of capturing the local network dynamics within the functionally relevant part of state space. Second, even when sampling locations are defined along the condition average trajectories, they need to be sampled at high enough density. When too few sampling locations are chosen along the trajectories, the identified global operative dimensions are less effective at reproducing network output and trajectories (Fig. S18).

One consequence of the tight link between local network dynamics and operative dimensions is that the number of operative dimensions required to approximate a particular network may increase with the complexity of the network activity, and of its inputs and outputs [23]. To illustrate this relation, we systematically varied the complexity of RNN computations by training sine wave generators with varying numbers of input and output signals (between 1 – 5 inputs *U* and outputs *Z*; Fig. 4). As expected, the network activity **x**_*t*_ becomes increasingly high-dimensional for increasing values of *U* and *Z* (Fig. 4b), and correspondingly the number of global operative dimensions required to achieve the original performance increases as well (Fig. 4c-e and f-h). We obtained a similar effect when increasing the dimensionality of the inputs into the RNNs, while keeping the dimensionality of the output fixed. (Fig. 13). Analytical considerations also imply that, in vanilla RNNs described by Eq. 1, the number of global operative dimensions is tightly linked to the dimensionality of the network activity **R** (section A.2.4 and A.2.5).

**Figure 4:**
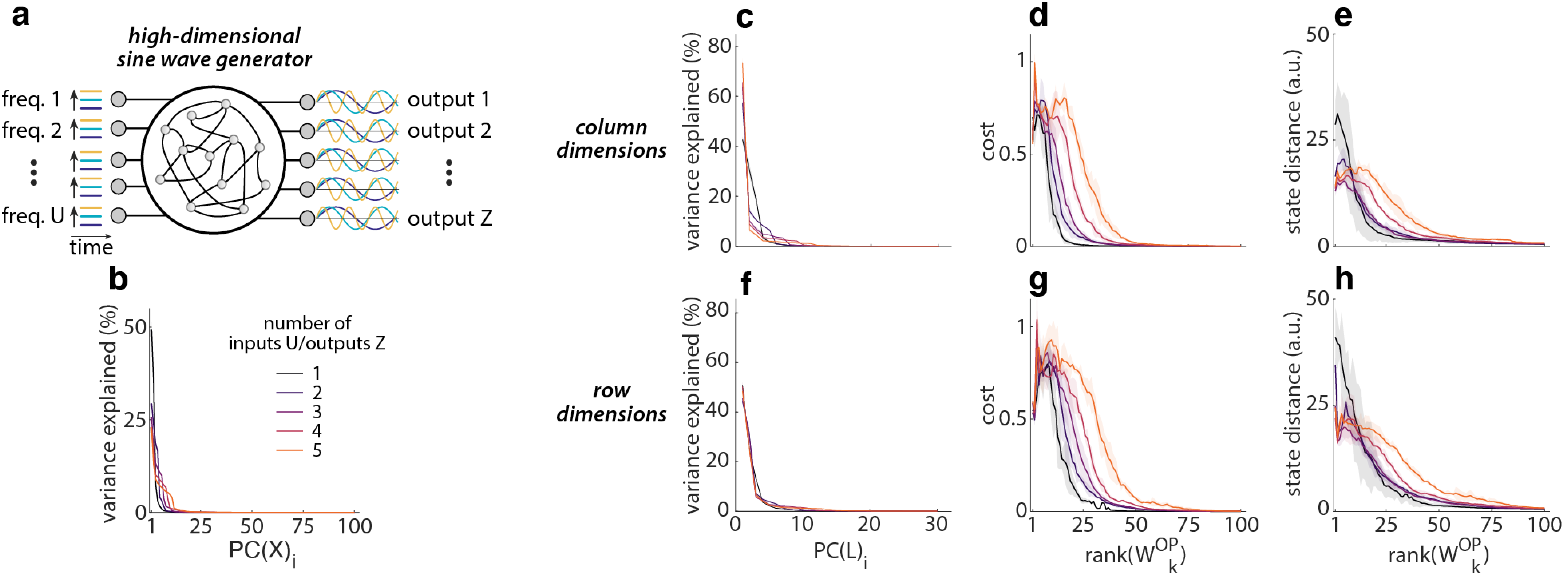
Operative dimensions and network complexity. (**a**) Task schematic for high-dimensional sine wave generator networks trained to output 1-5 sine waves simultaneously. (**b**) Variance explained (in activity space) by individual PCs of the network activity **X** over all input conditions. (**c**) Rank of global operative column dimensions, estimated with PC analysis on concatenated local operative dimensions **L** (Eq. 10 and 11). (**d**) Network output cost of networks with reduced-rank weight matrix 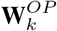 for *k* = 1 : *N* (Eq. 12). (**e**) State distance between trajectories in the full-rank network and in networks with reduced-rank weight matrix 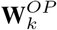. (**c-e**) Based on global operative *column* dimensions; averaged over 5 networks per number of inputs/outputs; shaded area: *mad*. Network output cost obtained with internal and input noise, state distance without any noise. (**f-h**) Same as (**c-e**) for global operative *row* dimensions.

#### 2.2.3 Operative dimensions relate functional modules to weight subspaces

Past studies have shown that RNNs can implement complex, context-dependent computations by “tiling” activity state space into separate functional modules [21, 19, 24]. Dynamics within individual modules is often approximately linear, but different approximately linear dynamics, corresponding to different input-output relations, are implemented across modules [20].

Our definition of operative dimensions is well-suited to ask whether the existence of functional modules in activity space has a correspondence at the level of the structure of the connectivity. To obtain a more fine-grained mapping from function to structure, the global operative dimensions can be generated based on different subsets of local operative dimensions. Specifically, sampling locations can be grouped based on their functional meaning, i.e. based on which condition average they belong to. The resulting sets of function-specific global operative dimensions can then link different network functions to particular subspaces in the weight matrix.

We demonstrate this approach for the context-dependent integrator network. We inferred global operative dimensions from local operative dimensions that were collected either separately by context, but pooled over choice, or separately by choice, but pooled over context (details section A.2.6). We refer to the resulting context-dependent global operative column dimensions as **q**_*i*_(*context_j_*), and the choice-dependent dimensions as **q**_*i*_ (*choice_j_*) (*i* = 1 : *N*, *j* = 1 : 2; row dimensions: 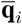). We compared these dimensions directly by measuring their pairwise alignment (subspace angle, Fig. 5a-d). We find that the function-specific global operative dimensions are only weakly aligned across the two contexts, but more strongly aligned across choices, suggesting that the RNN uses different weight subspaces to implement the sensory integration in each context, but reuses weight structures to implement the sensory integration across choices.

**Figure 5:**
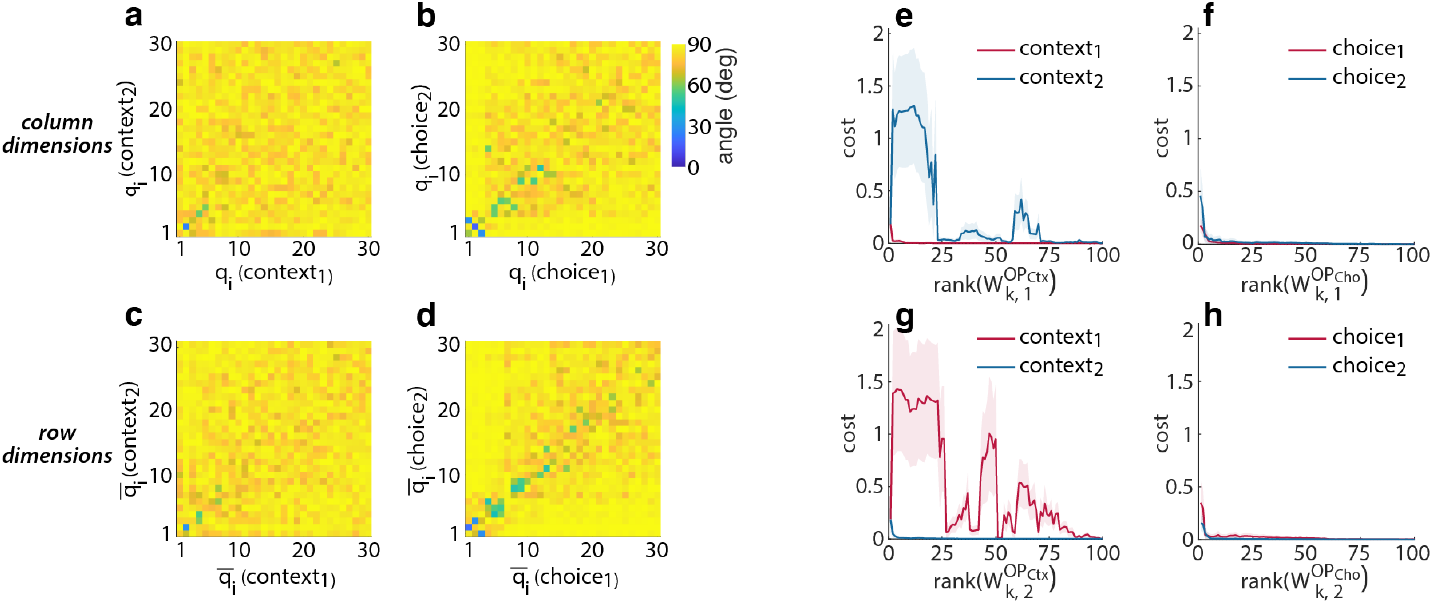
Function-specific global operative dimensions. (**a**) Subspace angle between global operative column dimensions **q**_*i*_ defined separately for context_1_ and context_2_. (**b**) Subspace angle between global operative column dimensions **q**_*i*_ defined separately for choice_1_ and choice_2_. (**c-d**) Same as (**a-b**) for global operative *row* dimensions 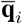. (**e**) Network output cost averaged separately over trials of context_1_ or context_2_ for networks with reduced-rank weight matrix consisting of the first *k* function-specific global operative column dimensions from context_1_ (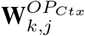, context *j* = 1). (**f**) Network output cost averaged separately over trials of choice_1_ or choice_2_ for networks with reduced-rank weight matrix consisting of the first *k* function-specific global operative column dimensions from choice_1_ (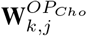, choice *j* = 1). (**g**) Same as (**e**), but based on global operative column dimensions from context 2. (**h**) Same as (**f**), but based on global operative column dimensions from choice 2.

To further support this conclusion, we checked if the global operative dimensions inferred from one functional module can support accurate computations in another module. We first constructed module specific, reduced-rank approximations of **W** as above (Eq. 12), which we refer to as 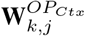 (contextspecific) and 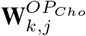 (choice-specific). We then measured network performance across conditions when using these different reduced-rank approximations (Fig. 5e-h). In agreement with the subspace-angles above, we find that networks based on operative dimensions defined in context1 perform poorly in context2, and vice versa (Fig. 5e, g) whereas operative dimensions from a given choice yield comparatively good performance also in the other choice (Fig. 5f, h). These observations are in line with previous findings [19] that the context-dependent integrator implements an approximate line attractor in each context, with mostly preserved dynamics along a given line attractor (i.e. across choices), but rather different dynamics between attractors (i.e. across context). Interestingly, the identified structure in the underlying connectivity, while clearly revealed by the operative dimensions (Fig. 5), does not become apparent when studying the weight matrix or contributions of individual network units directly (Fig. S19).

## 3 Discussion

In this work, we present an approach for inferring the *operative dimensions* of an arbitrary weight matrix in an RNN. The operative dimensions of the connectivity are defined based on their impact on the computationally relevant, local recurrent dynamics of the network. We find that for the examined tasks, the operative dimensions span a low-dimensional subspace of the connectivity space that is sufficient to produce accurate outputs in a reduced-rank network.

### Relation to state of the art

Our framework of operative dimensions extends recent work based on RNNs that by construction are explicitly low-rank, which already showed that many tasks of relevance to neuroscience can be solved with RNNs that rely entirely on low-rank connectivity [17].Unlike in this previous work, the networks we analyzed have a connectivity that is not constrained–the dimensionality of the underlying weight matrix is high at initialization, and remains high throughout training (Fig. 1d, i). This high dimensionality persists even though the weight updates brought about by training are low-rank [22]. However, our analyses showed that only a low-dimensional subspace of the effectively high-dimensional connectivity is used to solve the task, whereby the remainder of the structure in the weight matrix plays essentially no role in solving the task at hand. This finding opens the possibility to apply the conceptual framework of low-rank networks [17], which has provided valuable and direct insights into the relation of connectivity and dynamics, also to other types of RNNs.

Operative dimensions amount to a form of model reduction [25, 26] that emphasizes the preservation of attractor dynamics [27, 28] for the special case of a model parameterized as a neural network. Critically, our approach to relating the model structure to its function relies on characterizing the dynamics locally, along explored activity trajectories, similarly to methods to quantify non-linear dynamics based on Lyapunov or bred vectors [29, 30, 31].

More generally, our work makes a contribution towards increasing the *interpretability* of artificial neural networks. Past work had shown that unconstrained, trained RNNs need not be considered as a “black box”, but rather that the computations implemented by many RNNs can be understood at the level of population dynamics, through the interaction of inputs and dynamical primitives like attractors and saddle points [20, 32, 24]. Recent work related the implemented dynamical primitives to the underlying connectivity [17] in constrained RNNs. We have shown that establishing such relations is in principle also possible through the definition of condition-dependent operative dimensions (Fig. 5) without any specific constraints on the connectivity. Current opinions assume that such increased understanding and interpretability of artificial networks is desirable both to increase acceptance of the resulting machine learning applications throughout society, but also potentially to design better artificial system that overcome biases and limitations resulting from current approaches (see [33, 34, 35] for reviews). Nonetheless, potentially negative societal impacts of increased interpretability cannot be ruled out. For instance, if used on RNNs trained on personal data, operative dimensions may potentially facilitate the extraction of private information that was otherwise hidden in high-dimensional weight matrices.

### Limitations

One limitation of our approach to identifying operative dimensions is that it relies on studying local recurrent dynamics based on a somewhat arbitrary definition of sampling locations in activity state space. By design, the inferred operative dimensions will be sufficient to reproduce the full-rank dynamics only at these specific locations in state space. Note that there is no explicit requirement for the operative dimensions to reproduce the desired network output. We picked sampling locations along the condition average trajectories, based on the assumption that these average trajectories provide a good coverage of all the relevant local dynamics. In practice, this assumption may not hold in all RNNs. For one, average trajectories may travel through state-space regions that are not visited by any single trials, leading one to under-sample functionally relevant regions. For another, single-trial and average trajectories may also reflect dynamics that is not functionally relevant, and thus lead to over-sampling of regions that are not directly involved in generating the output. Further work may address alternative approaches for choosing sampling locations to optimize their functional relevance. To some extent, the adequacy of sampling locations can be tested empirically by systematically varying the number and location of sampling locations to optimize the performance of the inferred reduced-rank networks (Fig. S18).

Another potential limitation of our approach is that it may not be equally effective in identifying a functional subspace of the connectivity across all types of RNNs and in more complex tasks. While here we have focused on vanilla RNNs, we found that a low-dimensional functional subspace can be identified in such RNNs for a variety of network architectures, including with non-overlapping populations of excitatory and inhibitory neurons, or the use of different non-linear activation functions (Fig. S15). The properties of learned dynamics in the kind of tasks we employed is largely conserved across different types of networks (GRU, LSTM; [21, 24]), in particular the role of “tiling” activity space into distinct computational modules. This observation implies that our approach to relating connectivity and function at the level of local dynamics is at least meaningfully applicable even in these different kinds of networks. While we find that operative dimensions can retrieve a functionally relevant subspace also for a richer task like sequential MNIST (Fig. S16), it remains an open question whether these approaches will extend to harder AI problems.

### Conclusion

Operative dimensions can infer a functionally relevant subspace within highdimensional RNN connectivity that is sufficient to perform the task at hand. On one hand, this observation might benefit practical applications of RNNs as a computational tool, e.g. for continual learning in RNNs, by specifically protecting functionally relevant subspaces of the weight matrix to avoid catastrophic forgetting [36], or for weight compression, by storing the weight matrix as a linear combination of the global operative dimensions [37, 22]. On the other hand, the ability to identify functionally specific subspaces in the network connectivity may improve the applicability of RNNs as a model in neuroscience, as it simplifies the critical challenge of linking the properties of the connectivity to the network dynamics and may thus provide guidance on how to relate structure to function in complex biological datasets.

## Code Availability

We used Matlab to perform the data analysis. All the code and models required to reproduce the main analyses supporting our conclusions are available in matlab and python at: https://gitlab.com/neuroinf/operativeDimensions

## Author Contributions

R.K. and V.M. conceived the idea for the study, R.K. performed the analyses. M.C. and G.I. provided feedback throughout the project. R.K. and V.M. wrote the manuscript. R.K., V.M., M.C. and G.I. reviewed the manuscript.

## Acknowledgements

We thank all group members (Neural Computation and Cognition (V.M.), Cortical Computation (M.C.) and Neuromorphic Cognitive Systems (G.I.)) for their valuable feedback and discussions.

## A Appendix

### A.1 Recurrent neural networks

#### A.1.1 Task structures

##### Context-dependent integration

The context-dependent integrator [19] receives four inputs: two time-varying, noisy sensory inputs of six different levels, drawn at each time from 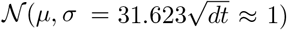 with *μ* ∈ {–0.5, –0.12, –0.3, 0.03, 0.12, 0.5}; and two constant context inputs, set to 1 or 0. In context_1_, input context_1_ is set to 1 and input context_2_ is set to 0 while both sensory inputs are ON. The task is to integrate the relevant sensory input (input sensory_1_) over time in the activity of the output unit **z**_*t*_ and ignore the irrelevant sensory input (input sensory_2_) and vice versa for context_2_. Each trial consists of 650 *ms* burn period (only input context_j_=1; both sensory inputs OFF) followed by 750 *ms* sensory integration time (only input context_j_=1; both sensory inputs ON). One trial has *T* = 1400 time steps.

##### Sine wave generation

The sine wave generator [20] receives one constant input which can be set to one of 51 different levels 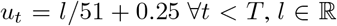. The level of the constant input defines the target frequency of the sine wave generated in the activity of the network output unit **z**_*t*_ whereby the target frequencies *ω_l_*(*l* = 1 : 51) are equally spaced between 1 – 6 *rad/sec* to define *ω_l_* = 0.1 * (*l* – 1) + 1. Each trial lasts 500 *ms* during which the input *u_t_* is constantly ON (*T* = 500).

##### High-dimensional sine wave generation

The high-dimensional sine wave generator network (Fig. 4) is an extension of the sine wave generator network explained in section A.1.1 [20]. It receives *U* =1 : 5 constant inputs **u**_*t*_ and is trained to generate sine wave activity in *U* = *Z* =1 : 5 outputs **z**_*t*_. The *U* constant inputs are set independently of each other to one of 51 different levels 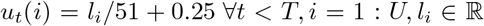. The level of the i-th constant input *u_t_*(*i*) defines the target frequency of the sine wave generated in the activity of the i-th network output unit *z_t_*(*i*) whereby the target frequencies *ω_l_i__*(*l_i_* = 1 : 51) are equally spaced between 1 – 6 *rad/sec* to define *ω_l_i__* = 0.1 * (*l_i_* – 1) + 1. Each trial lasts 500 *ms* during which the input is constantly ON (*T* = 500). For *U* = *Z* = 1 the high-dimensional sine wave generation network is identical to the sine wave generation network explained in section A.1.1. We train separate networks for every number of *U* = *Z* =1 : 5.

#### A.1.2 Network training

In both tasks, all weights (**B**, **W**, **Y**) are optimized using Hessian-free optimization [38] to minimize the network output cost:

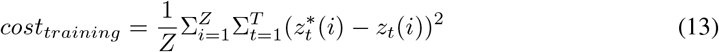

between the respective target activity 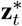 and the network output activity **z**_*t*_ at every time point during the trial, averaged over all input conditions (figures show *cost_training_* scaled by 1/*T*, Eq. 3). **B**, **W**, and **Y** are randomly initialized and set to have a spectral radius of 1 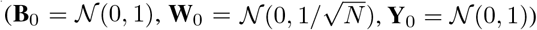. All input conditions are trained simultaneously with equal probability to ensure a balanced training set (batch size = 400). The presented results are obtained using the same input conditions as during training but with varying input noise and internal noise. The code to train and run the RNNs was modified from [19]. For the main results we trained 20 networks on contextdependent integration, 20 networks on sine wave generation and 25 networks on high-dimensional sine wave generation (5 networks per *U* = *Z* (*U* = 1 : 5, *Z* = 1 : 5)).

### A.2 Definition of operative dimensions

#### A.2.1 Sampling locations

We place the sampling locations **y**_*j*_ (*j* = 1 : *P*) equally spaced in time on the condition average trajectories of each network. The condition average trajectories are defined as the mean network activity per time point of the corresponding input condition, averaged over input and internal noise (20 trials per condition).

For the context-dependent integration network we considered the condition average trajectories for 8 different input conditions whereby we define the input condition based on the context (1 or 2), the choice (1 or 2) and the coherency of the sensory inputs (congruent (sign(input sensory_1_) = sign(input sensory_2_) or incongruent (sign(input sensory_1_) != sign(input sensory_2_)) in every trial (2^3^ = 8 distinct input conditions). We placed the sampling locations equally spaced in time along each of these 8 condition average trajectories at every 100-th time step (t=1:100:1400 resulting in 15 sampling locations per condition average trajectory) to obtain **y**_1_ = **x**_1_, **y**_2_ = **x**_100_, **y**_3_ = **x**_200_, … for every input condition. In total, we defined *P* = 8 × 15 = 120 sampling locations.

The sine wave generation network has 51 different input conditions (51 input levels). For the definition of local operative dimensions, we subsampled the input conditions and only considered every 5-th input level (*l* = 1 : 5 : 51), resulting in 11 condition average trajectories. We placed the sampling locations equally spaced in time along each of these 11 condition average trajectories at every 50-th time step (t=1:50:500 resulting in 11 sampling locations per condition average trajectory) to obtain **y**_1_ = **x**_1_, **y**_2_ = **x**_50_, **y**_3_ = **x**_100_, … for every input condition. In total, we defined 11 × 11 = 121 sampling locations.

#### A.2.2 Operative row dimensions

Analogously to the operative *column* dimensions, the operative *row* dimensions are defined as the dimensions in **W** that have a large impact on the local dynamics if removed from the *row* space of **W** (Fig. 3a).

Given an arbitrary unit vector 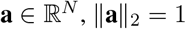, we define 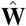 for the operative *row* dimensions as the matrix of rank *N* – 1 obtained by removing the dimension **a** from the *row* space of **W**:

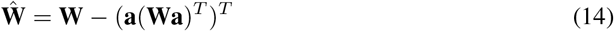

The respective local and global operative dimensions are defined as explained in section 2.2.1. Based on Eq. 7, 8, 9, 10, 11, the i-th global operative row dimension is defined as the i-th left singular vector of **L** and we refer to is as 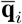.

The reduced-rank approximation of **W** constructed from only a subset of the global operative *row* dimensions is then given by:

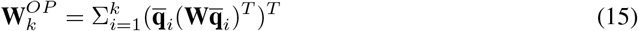

For simplicity and readability purposes, below we use the variables for the global operative *column* dimensions throughout the text. All corresponding statements similarly apply also to the row dimensions, unless explicitly stated otherwise.

#### A.2.3 Properties of local operative dimensions in vanilla RNNs

For the special case of a vanilla RNN (Eq. 1), several properties of the local and global operative dimensions can be derived analytically. While the derivations below do not apply to other RNN architectures (e.g. LSTM [39], GRU [40]) the general approach and definitions for the estimation of operative dimensions are applicable irrespective of the RNN architecture.

##### Analytical derivation of local operative column dimensions

The first local operative dimension **d**_1, *j*_ at location **y**_*j*_ is the solution of the following optimization problem (Eq. 8):

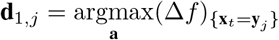

with 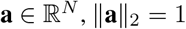, and Δ*f* as (Eq. 1, 7):

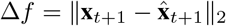

For a vanilla RNN:

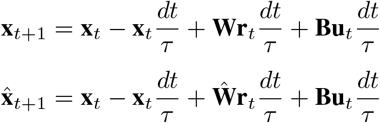

To derive the first local operative *column* dimension this can be simplified as follows:

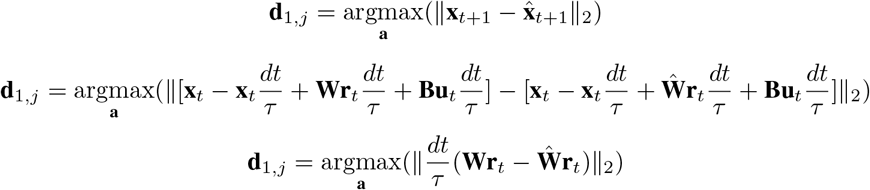

We replace 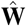 using Eq. 6:

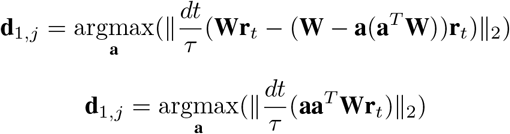

which has a unique solution for a vector **a** that is aligned to **Wr**_*t*_:

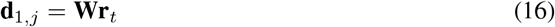

##### Dimensionality of local operative column dimensions

From the above derivation also follows that, at any given location **x**_*t*_ in the state space of a vanilla RNN, only a single local operative column dimension can be derived. This follows from the observation that removal from **W** of any vector that is orthogonal to **d**_1, *j*_ = **Wr**_*t*_ necessarily results in Δ*f* = 0, and thus does not cause any change in the network dynamics:

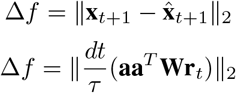

Replacing **Wr**_*t*_ = **d**_1, *j*_:

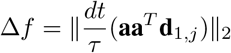

Given that all local operative column dimensions **d**_*i,j*_ ∀*i* > 1 have to fulfill 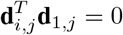 (see Eq. 9), it follows that Δ*f*_*i,j*_ =0, ∀*i* > 1

Hence, when using the standard RNN equation as described in Eq. 1, only a single local operative *column* dimensions **d**_1, *j*_ = **Wr**_*t*_ can be inferred at any given location in state space.

The previous two key derivations on local operative column dimensions can further be extended to the case where the non-linear RNN dynamics is well described by the local linear approximation 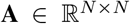 around slow points of the full-rank RNN 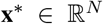. Hence, the derivations and corresponding results are more general and apply (approximately) to any RNN architecture that results in dynamics that are locally linear, not just to vanilla RNN.

#### A.2.4 Dimensionality of global operative *column* dimensions

The above properties of the *local* operative column dimensions have implications for the overall dimensionality of the *global* operative column dimensions. Specifically, in a vanilla RNN the dimensionality of the subspace spanned by the global operative column dimensions is bounded by the dimensionality of the network activity **r**_*t*_. This follows from the above derivation (section A.2.3) that there is only a single local operative column dimension at any sampling location **y**_*j*_ in state space, namely **d**_1, *j*_ = **Wr**_*t*_ with **r**_*t*_ = *tanh*(**y**_*j*_).

The population activity **r**_*t*_ at a given location in state space and time can be written as a linear combination of the principal components of the population activity *PC*(**R**)_*i*_:

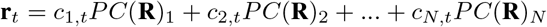

where the coefficients *c_i,t_* (*i* = 1 : *N*) depend on state space location and time *t* in the trial. Combining the above expression with **d**_1, *j*_ = **Wr**_*t*_ results in:

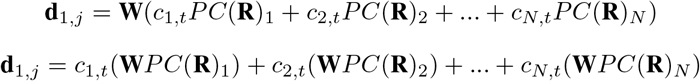

While the coefficients *c_i,t_* vary over sampling locations **r**_*t*_ = *tanh*(**y**_*j*_), the vectors **W***PC*(**R**)_1_, **W***PC*(**R**)_2_, …, **W***PC*(**R**)_*N*_ do not, and rather are a fixed property of an RNN with weight matrix **W**. Hence, all local operative column dimensions are a linear combination of the **W***PC*(**R**)_*i*_ with *c_i,t_* > 0. As a consequence, if the population activity **R** is contained in a low-dimensional subspace, the subspace of the column space of **W** that is required to perform the task (spanned by the operative column dimensions) is also low-dimensional, and its dimensionality is at most as high as the dimensionality of the responses **R**, independently of the training procedure. Note that while the dimensionality of these two subspaces is related, the two subspaces need not be overlapping.

Notably, a comprehensive understanding of the exact factors determining the dimensionality of activity in trained RNNs is currently lacking. The dimensionality of the inputs can be expected to be an important factor [1] although in general not the only one. Indeed, RNNs that receive high-dimensional inputs can nonetheless generate low-dimensional dynamics [32]. On the other hand, reservoir computing networks can generate high-dimensional dynamics even when driven with low-dimensional inputs [41, 42]. Our result that the dimensionality of activity in the N-fold sine-wave generator increases with N (Fig. 4b) could further be interpreted as being driven by the dimensionality of the output.

#### A.2.5 Dimensionality of global operative *row* dimensions

Analytical constraints to the functional subspace spanned by the global operative *row* dimensions can also be derived. Specifically, this functional subspace is contained within the intersection of the subspace spanned by the network activity **R** and the subspace spanned by the row dimensions of **W**. Indeed, any vector orthogonal to this intersection can be removed from **W** without any effect on the network dynamics.

The effect of removing dimension **a** from the row space of **W** is given by:

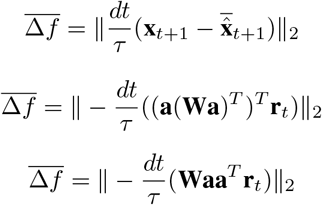

The term **Wa** vanishes for any **a** outside the row space of **W**. The term **a**^*T*^**r**_*t*_ vanishes for any term orthogonal to **r**_*t*_. Hence, 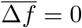 for any **a** outside the intersection between the row space of **W** and the activity subspace.

#### A.2.6 Function-specific global operative dimensions

To define function-specific global operative dimensions, we combine local operative dimensions only from specific subsets of sampling locations. In Fig. 5, we created four different types of such function-specific global operative dimensions **q**_*i*_(*function*) (*i* = 1 : *N*) for the context-dependent integration network:

- **q**_*i*_(*context*_1_): sampling locations *y_j_* along the condition average trajectories of context_1_
- **q**_*i*_(*context*_2_): sampling locations *y_j_* along the condition average trajectories of context_2_
- **q**_*i*_(*choice*_1_): sampling locations *y_j_* along the condition average trajectories of choice_1_
- **q**_*i*_(*choice*_2_): sampling locations *y_j_* along the condition average trajectories of choice_2_

All four types combine sampling locations from 4 condition average trajectories with 15 sampling locations each (*P* = 60).

#### A.2.7 Required computational resources

Computing the local operative column dimensions is computationally inexpensive because there is an explicit solution (**d**_1, *j*_ = **Wr**_*t*_; see section A.2.3) and hence it is not required to run the numerical optimization procedure but only 1 matrix multiplication per sampling location.

To obtain the local operative row dimensions we perform a numerical optimization (matlab function *fminunc* to *find the minimum of an unconstrained multivariable function* using the *quasi-newton* optimization algorithm; as provided in the code to obtain the operative dimensions). We run the optimization upto N-times per sampling location or until Δ*f* < 1*e*^−8^. It takes less than one minutes to obtain the local operative row dimensions at a given sampling location on a standard machine (6 core, Intel Core i7-7800X CPU, 3.50GHz).

### A.3 Additional analyses and figures

#### A.3.1 Importance of initial rank of W

The trained weight matrices **W** are generally high-dimensional in both tasks (Fig. 1d, i). Interestingly, the rank of the trained weight matrix seems highly dependent on the rank of the initial weight matrix **W**_0_ (Fig. S6) whereby a low-dimensional, initial weight matrix **W**_0_ generally results in a low-dimensional, trained weight matrix **W**.

#### A.3.2 High-variance dimensions

To assess the functional importance of the high-variance dimensions of **W**, we sequentially remove the high-variance dimensions from **W** while measuring the performance of the reduced-rank network (Fig. 1e, j). Generally, the network performance shows sudden jumps at specific ranks which are hard to interpret. Here we show how the network performance varies over reduced-rank 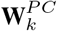 for more example networks to illustrate the large differences that are seen across individual networks (Fig. S7).

#### A.3.3 Alignment between global operative dimensions and high-variance dimensions

The large difference between operative and high-variance dimensions also becomes apparent when comparing their pairwise alignment to each other. While the subspace angles between the first few global operative dimensions to the first few high-variance dimensions show a weak alignment, the remaining dimensions are almost orthogonal to each other (Fig. S8; *subspace angle* = *acos*(|**q**_*i*_ · *PC*(**W**)_*j*_|)).

Interestingly, the global operative column dimensions show a stronger alignment with the linear network activity *PC*(**X**) (Fig. S8a, c) and the global operative row dimensions with the non-linear network activity *PC*(**R**) (Fig. S8b, d). However, despite their partial alignment the PCs of the network activity are not describing the functionally relevant subspace as accurately as the global operative dimensions. To assess this, we construct reduced-rank approximations of **W** using *PC*(**X**)_*i*_:

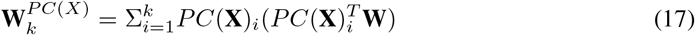

and similarly *PC*(**R**)_*i*_:

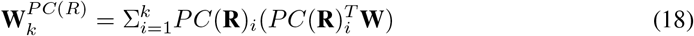

Analogously for row dimensions based on Eq. 15. The networks require a larger number of *PC*(**X**)_*i*_ or *PC*(**R**)_*i*_ than global operative dimensions in **W** to achieve the same performance level (Fig. S9).

#### A.3.4 Alignment to other relevant dimensions

While the operative dimensions are not well aligned with the high-variance dimensions shown above, they may be aligned with other important dimensions of the RNN. Here we consider three additional types of dimensions: (1) the input dimensions; (2) the eigenvectors of the weight matrix; and (3) the main dimensions characterizing the local linear dynamics around the chosen sampling locations.

First, we considered the alignment between the global operative dimensions and the input dimensions (Fig. S10), which we define based on various different approaches: we consider the principal components of the input weight matrix **B** (panel a; *subspace angle* = *acos*(|**q**_*i*_ · *PC*(**B**)_*j*_ |)); the main directions in state space effectively explored by the inputs (panel b; *subspace angle* = *acos*(|**q**_*i*_ · *PC*(**Bu**_*t*_)_*j*_ |) ∀*t* = 1 : *T*); or directly the column space of **B** (panel c; *subspace angle* = *acos*(|**q**_*i*_ · *column*(**B**)_*j*_)|) ∀*j* = 1 : *U*). Irrespective of the employed definition, all these inputs dimensions are largely orthogonal to the column operative dimensions, and only weakly aligned to a few row operative dimensions. Some alignment with the row operative dimensions can be expected, as these dimensions in **W** describe the *input* connections of the hidden units. It seems reasonable that the functionally relevant subspace of the row space in **W** - described by the global operative row dimensions - is at least partially aligned with the task input dimensions, as these mediate the inputs that are crucial drivers of the network activity while performing the task.

Second, we consider the alignment between the global operative dimensions and the right and left eigenvectors of **W** (Fig. S11; *subspace angle* = *acos*(|**q**_*i*_ · *right/left eigenvector* (**W**)_*j*_ |)). Both eigenvectors are at most weakly aligned to the global operative dimensions, emphasizing that our approach retrieves dimensions that may not be directly identifiable based on the weight matrix alone.

Third, we considered the alignment between a local operative dimensions estimated at a particular state-space location and the linearized RNN dynamics **A** at that location (linearized at sampling locations on condition average trajectory, see Eq. 24 and 25 for definition of **A**). In Fig. S12 we characterize the linear dynamics through the *PC* of of **A**, which are not aligned with with respective first local operative dimensions at any sampling location (large subspace angles; *subspace angle* = *acos*(|**d**_1, *j*_ · *PC*(**W**)_*j*_|)). Likewise, we failed to find any alignment between the local operative dimensions and the right and left eigenvectors of **A** at any sampling location (not shown).

The mismatch between operative dimensions and linear dynamics might at least partly reflect our definition of operative dimensions, which is based entirely on the contribution of the recurrent dynamics **Wr** while discarding the decay term −**x***dt*/*τ* (see Eq. 1). The linearized dynamics, on the other hand, includes contributions from both terms. Hence, an interesting extensions of our presented definition of operative dimensions would additionally consider the decay term to define 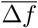 and remove **a** from the decay term similar to as from **W** (Eq. 6). However, such a formulation would make the resulting operative dimensions harder to interpret and more work is required to gain more insights into such alternative definitions of operative dimensions.

#### A.3.5 Importance of input dimensionality

To understand if the dimensionality of the global operative dimensions is related to the dimensionality of the network input we study operative dimensions in networks with systematically varied number of inputs. Specifically, we train RNNs (*N* = 100) on an extended version of the context-dependent integration task in which the networks are trained to distinguish between up to 9 contexts simultaneously (*U* = 1 : 9, *Z* = 1, Fig. 13a). We trained 5 networks for every *U* (*U* = 1 : 9, *Z* = 1) and obtained the global operative dimensions for each of them.

**Figure 6:**
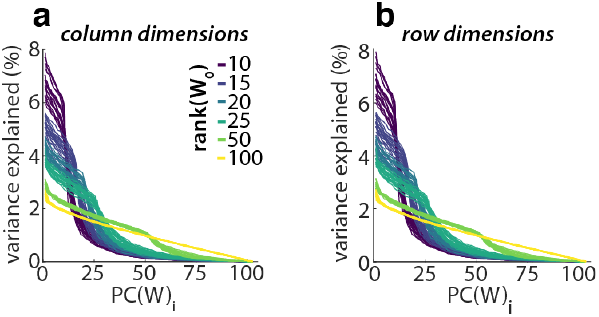
Dimensionality of trained weight matrices with different initial ranks. (**a**) Variance explained (in weight space) by individual PCs of the weight matrix **W** after training when the initial weight matrix was randomly initialized with different ranks (colors, see legend). (**b**) Same as (**a**) for row dimensions of **W**. (**a** - **b**) Each line is one network; 20 networks per **W**_0_; trained on context-dependent integration. For the column as well as the row space, the rank of **W** changes little over training, and is instead mainly determined by the rank of **W**_0_. The dimensionality of the weight matrix **W** after training is thus only weakly related to the trained task in these cases.

**Figure 7:**
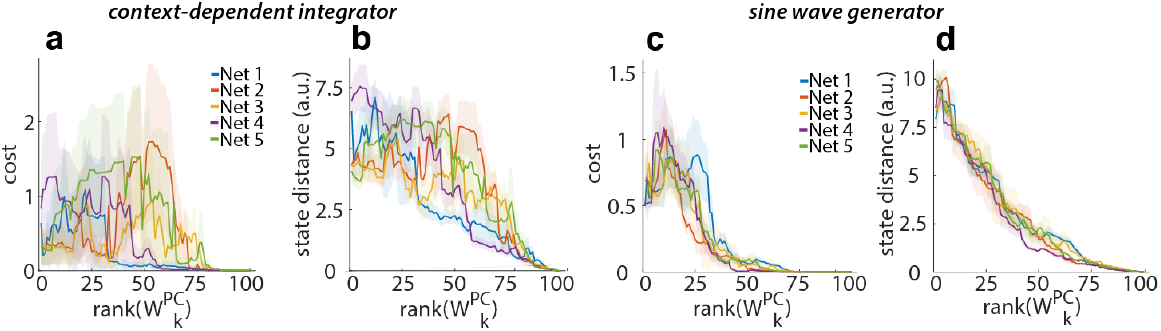
Network performance for reduced-rank weight-matrices based on high-variance dimensions. (**a**) Network output cost (Eq. 3) of networks with reduced-rank weight matrices 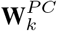 for *k* = 1 : *N* (Eq. 5). (**b**) State distance between trajectories in the full-rank network and in networks with reduced-rank weight matrix 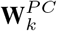. (**c-d**) Same as (**a-b**) for 5 example networks trained on sine wave generation. (**a-d**) Shown for 5 networks each; shaded area: *mad* over trials; Network output cost obtained with internal and input noise, state distance without any noise. The network performance shows large jumps at specific ranks that differ across networks (similar to Fig. 1 e, j).

**Figure 8:**
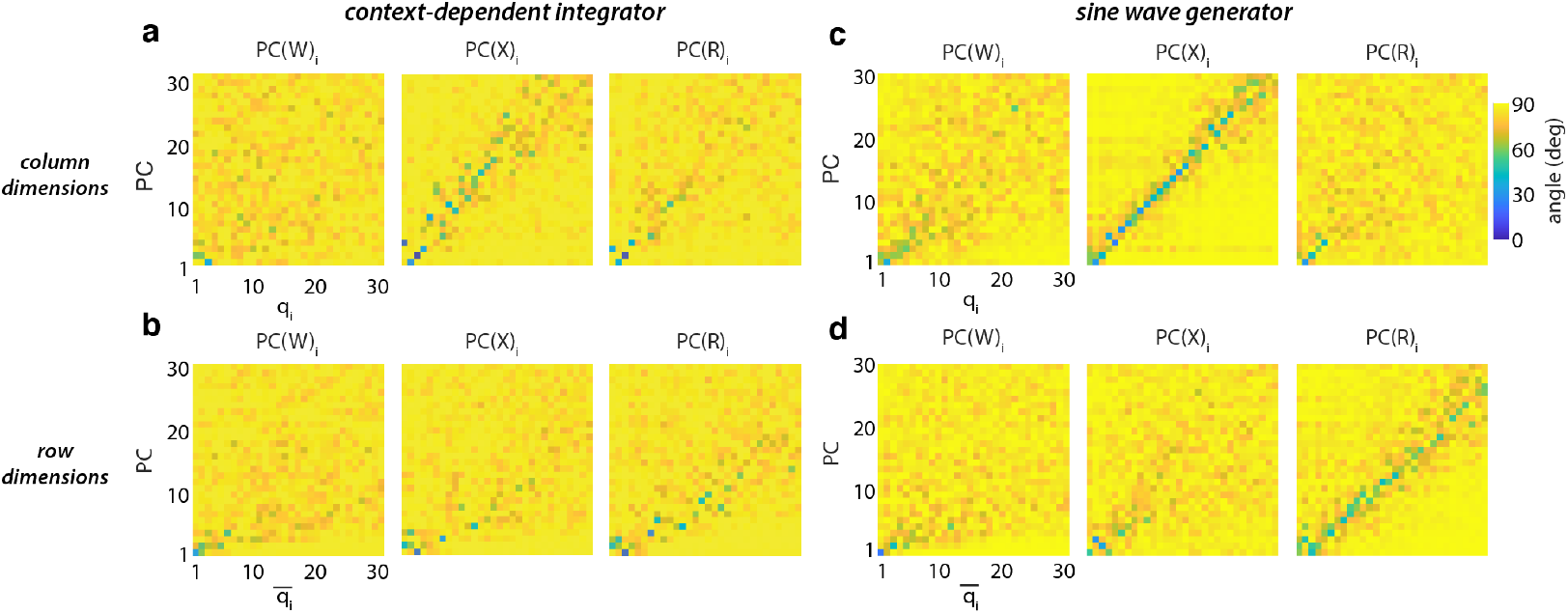
Alignment of operative to high-variance dimensions. (**a**) Subspace angle between global operative column dimensions **q**_*i*_ and PCs of **W**, **X** and **R**. (**b**) Same as (**a**) for global operative *row* dimensions in context-dependent integration. (**c**) Same as (**a**) for global operative *column* dimensions in sine wave generation. (**d**) Same as (**a**) for global operative *row* dimensions in sine wave generation. (**a** - **d**) Average over 20 networks per task.

**Figure 9:**
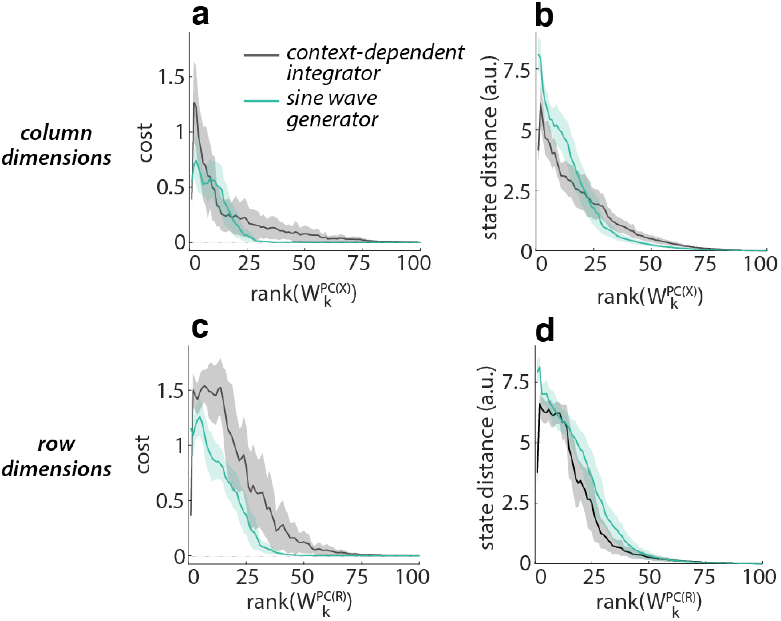
Network performance for reduced-rank RNN based on PCs of network activity X and R. (**a**) Network output cost of networks with reduced-rank weight matrix 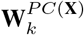 for *k* = 1 : *N* (Eq. 17). (**b**) State distance between trajectories in the full-rank network and in networks with reduced-rank weight matrix 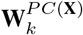 for *k* = 1 : *N*. (**c** - **d**) Same as (**a** - **b**) for removing *PC*(**R**)_*i*_ from row dimensions of **W**. (**a** - **d**) Averaged over 20 networks per task; shaded area: *mad* over networks; Network output cost obtained with internal and input noise, state distance without any noise.

**Figure 10:**
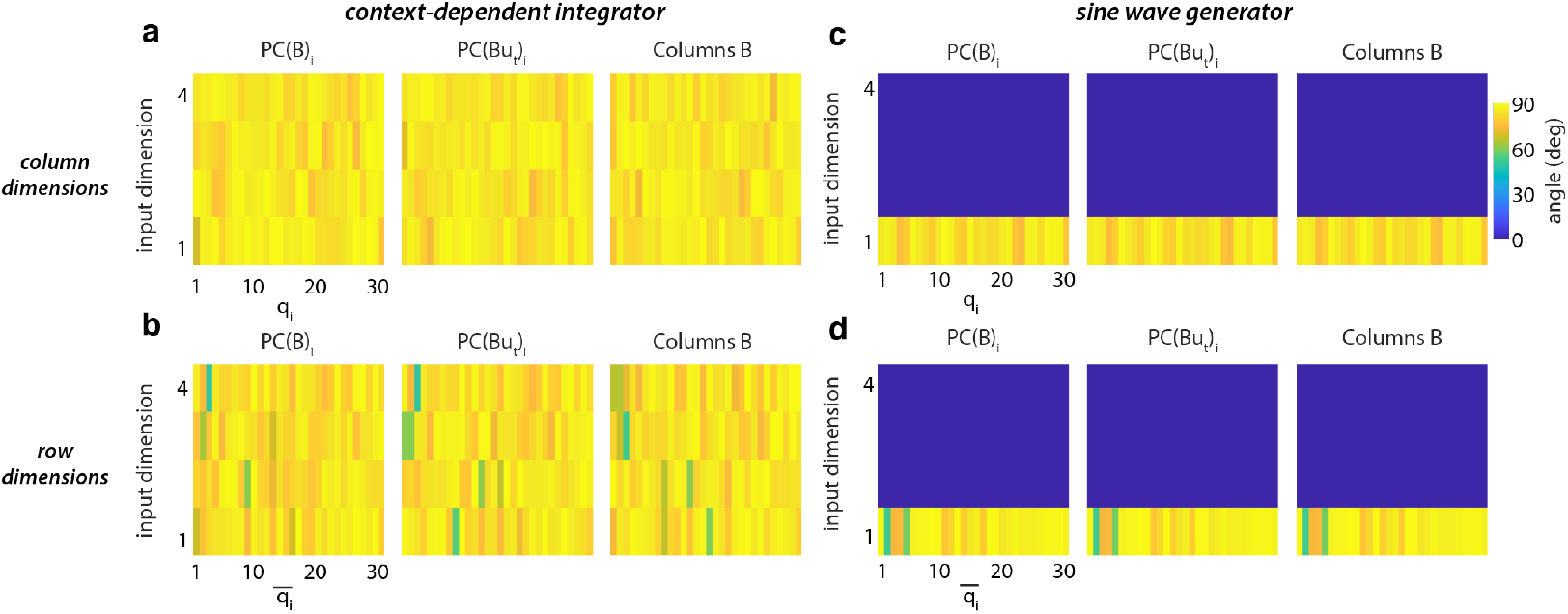
Alignment between global operative dimensions and network input dimensions. (**a**) Subspace angle between global operative column dimensions **q**_*i*_ and PCs of **B**, the PCs of **Bu**_*t*_, and the columns of **B**. (**b**) Same as (**a**) for global operative *row* dimensions in context-dependent integration. (**c**) Same as (**a**) for global operative *column* dimensions in sine wave generation. (**d**) Same as (**a**) for global operative *row* dimensions in sine wave generation. (**a** - **d**) Average over 20 networks per task.

**Figure 11:**
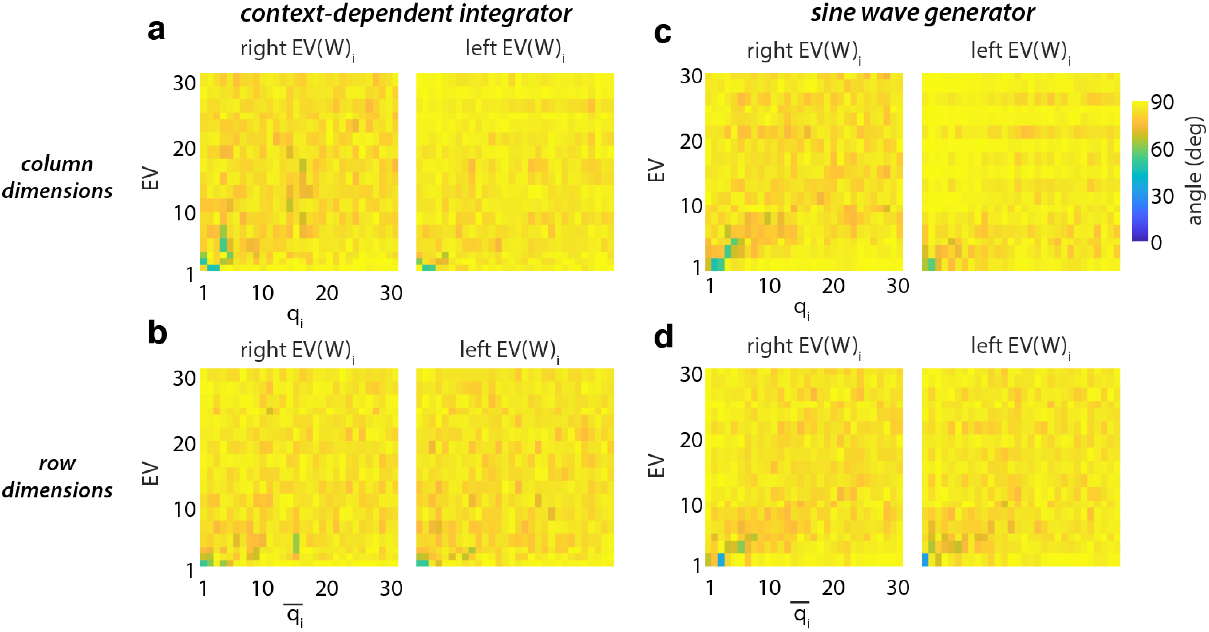
Alignment between global operative dimensions and eigenvectors of W. (**a**) Subspace angle between global operative column dimensions **q**_*i*_ and right and left eigenvectors of **W**. (**b**) Same as (**a**) for global operative *row* dimensions in context-dependent integration. (**c**) Same as (**a**) for global operative *column* dimensions in sine wave generation. (**d**) Same as (**a**) for global operative *row* dimensions in sine wave generation. (**a** - **d**) Average over 20 networks per task.

**Figure 12:**
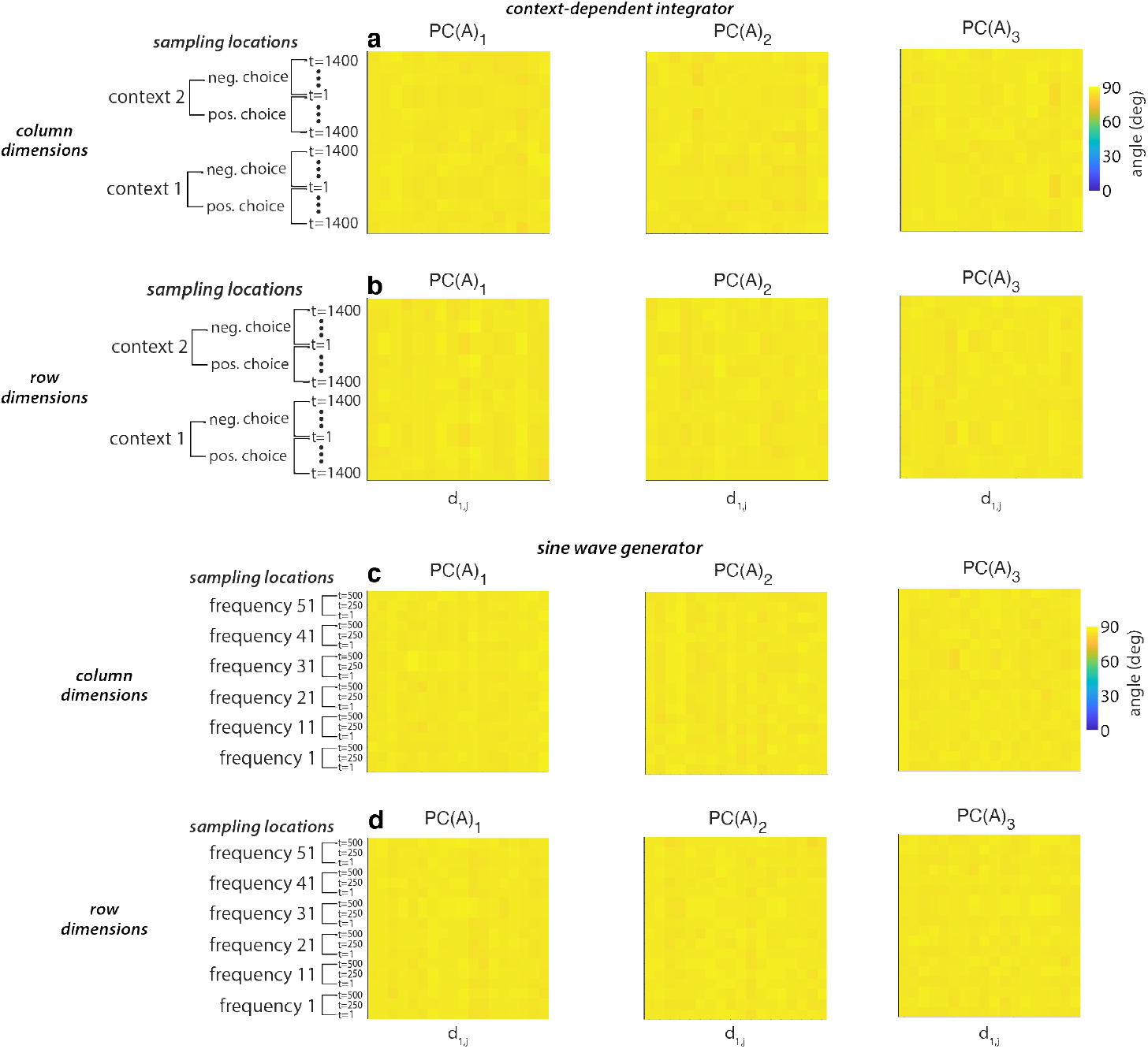
Alignment between local operative dimensions and linearized dynamics. ***A***. (**a**) Subspace angle between the first local operative column dimensions **d**_1, *j*_ and the first three PCs of **A** compared over sampling locations. (**b**) Same as (**a**) for global operative *row* dimensions in context-dependent integration. (**c**) Same as (**a**) for global operative *column* dimensions in sine wave generation. (**d**) Same as (**a**) for global operative *row* dimensions in sine wave generation. (**a** - **d**) Average over 20 networks per task.

**Figure 13:**
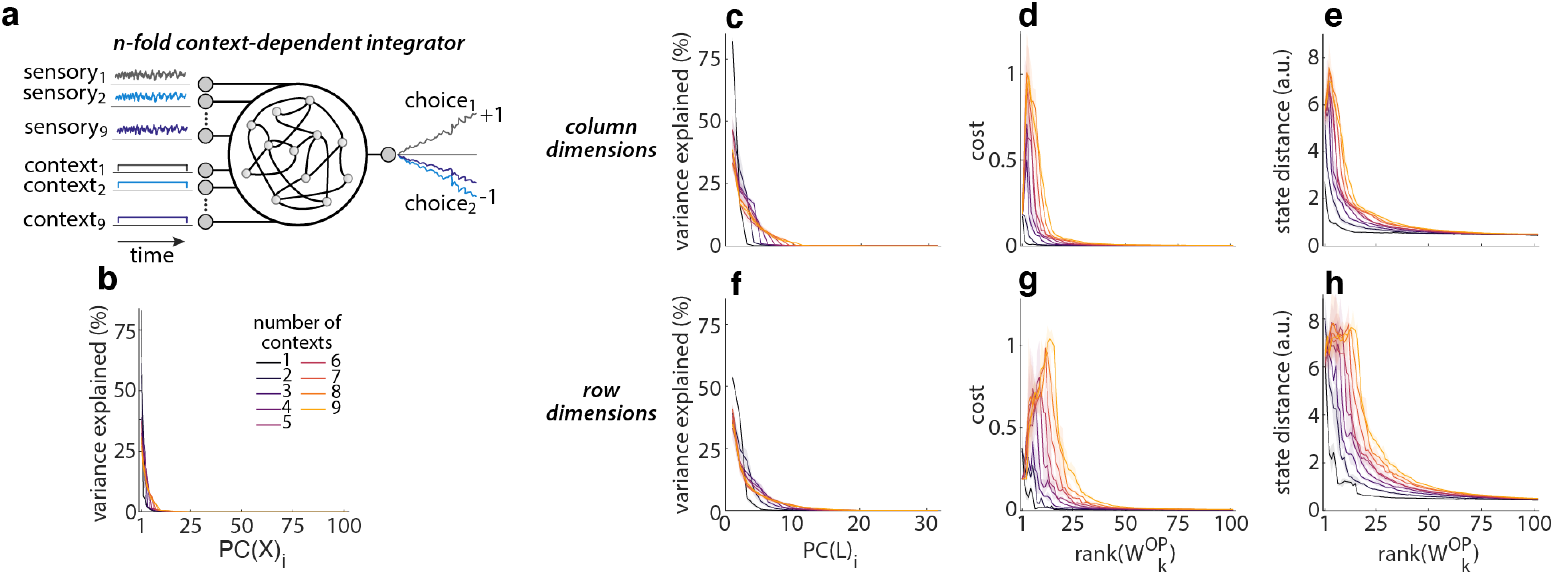
Operative dimensions and input dimensionality. (**a**) Task schematic for n-fold contextdependent integrator networks trained to select and integrate the relevant input out of 1-9 sensory inputs simultaneously. (**b**) Variance explained (in activity space) by individual PCs of the network activity **X** over all input conditions. (**c**) Rank of global operative column dimensions, estimated with PC analysis on concatenated local operative dimensions **L** (Eq. 10 and 11). (**d**) Network output cost of networks with reduced-rank weight matrix 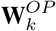 for *k* = 1 : *N* (Eq. 12). (**e**) State distance between trajectories in the full-rank network and in networks with reduced-rank weight matrix 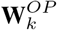. (**c-e**) Based on global operative *column* dimensions; averaged over 5 networks per number of contexts; shaded area: *mad*. Network output cost obtained with internal and input noise, state distance without any noise. (**f-h**) Same as (**c-e**) for global operative *row* dimensions.

In networks with higher number of inputs the dimensionality of the global operative dimensions is generally higher (Fig. 13c, f) and they require a larger number of dimensions to perform the task with the same performance (Fig. 13d,e,g,h). The overall dimensionality of the network activities also increases with *U* (Fig. 13b) which further demonstrates the tight link between the dimensionality of the network activity and operative dimensions. Note that the dimensionality of **W** remains roughly the same over all *U* (not shown).

#### A.3.6 Operative dimensions over training

Operative dimensions can be estimated for the RNN at any stage of training. The global operative dimensions at a particular training stage are defined based on sampling locations **y**_*j*_ located along the condition average trajectories for that training stage. Sampling locations are placed along the trajectories equally spaced in time, as described in section 2.2.1. For all training stages, a lowdimensional subspace in **W** can be defined which is sufficient to achieve the performance of the full-rank network at the same training stage (Fig. 14). Note that the network output cost for the full-rank networks is higher at early stages in training.

**Figure 14:**
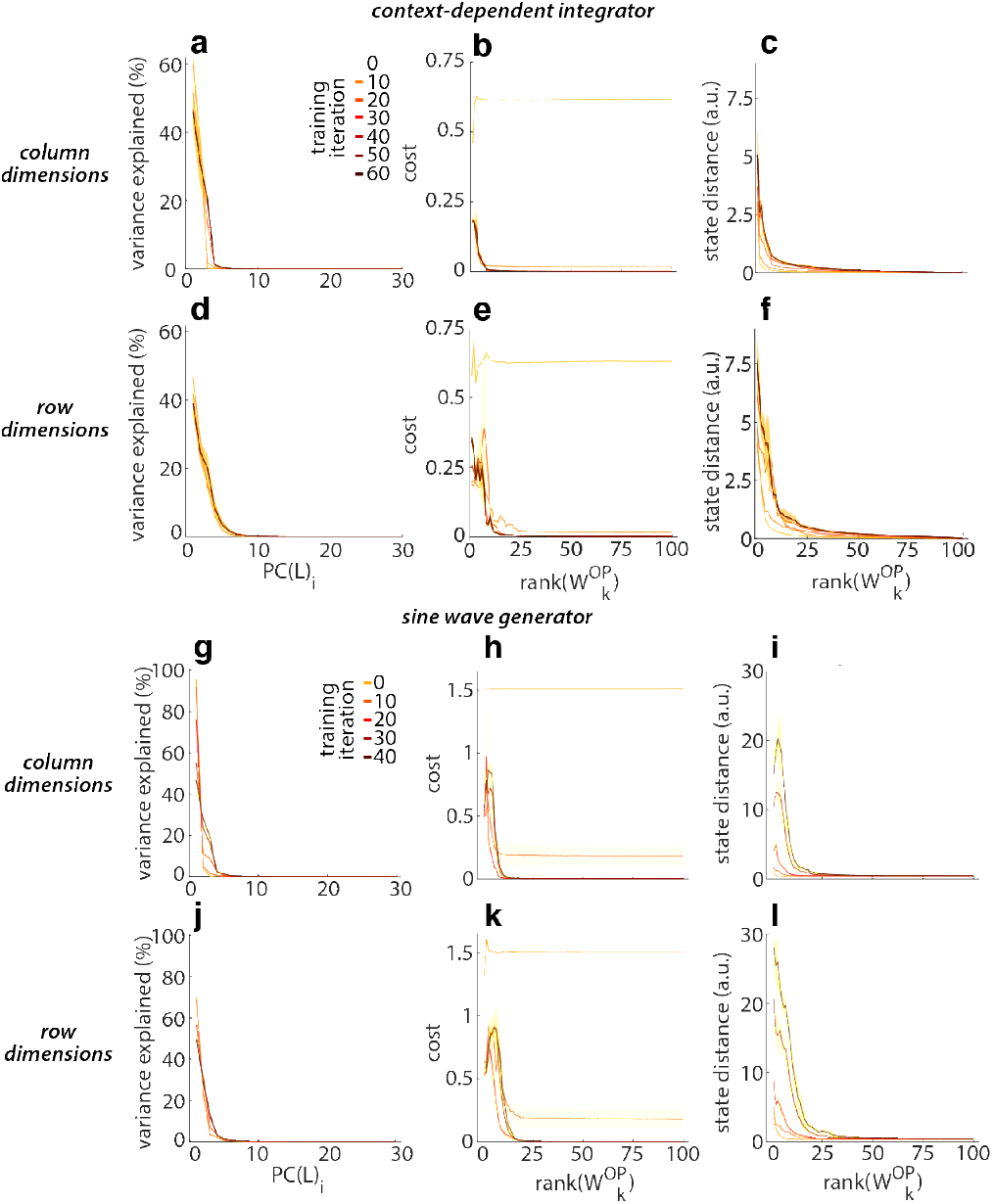
Operative dimensions over training. (**a**) Rank of global operative column dimensions, estimated with PC analysis on concatenated local operative dimensions (Eq. 10 and 11) for networks trained on context-dependent integration at different stages of training. (**b**) Network output cost of networks with reduced-rank weight matrix 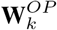 for *k* = 1 : *N* (Eq. 12) trained on context-dependent integration at different stages of training. (**c**) State distance between trajectories in the full-rank network and in networks with reduced-rank weight matrix 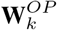 trained on context-dependent integration at different stages of training. (**d-f**) Same as (**a-c**) for global operative *row* dimensions in context-dependent integration networks. (**g-i**) Same as (**a-c**) for global operative *column* dimensions in sine wave generation networks. (**j-l**) Same as (**a-c**) for global operative *row* dimensions in sine wave generation networks. (**a-l**) Averaged over 5 networks per training iteration; shaded area: *mad* over networks; Network output cost obtained with internal and input noise, state distance without any noise.

#### A.3.7 Operative dimensions for different network types

To illustrate the general applicability of our definition of operative dimensions, we also estimated operative dimensions for the following three alternative types of RNNs:

##### bias, x0

We extended the standard RNN equation (Eq. 1) with trainable parameters for bias of hidden units 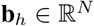, bias for output units 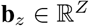 and initial conditions per context_j_ 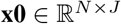 for *J* = 2 in the context-dependent integration network.

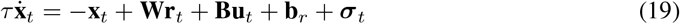

where **x**_*t*=0_ is set to **x0**_*j*_ for trials of context_j_. The network output is defined as:

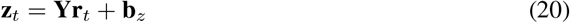

##### relu, bias, x0

Same as Eq. 19 and 20, but replacing the *tanh* with the *relu* activation function for **r**_*t*_:

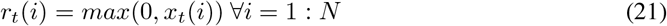

##### dale’s law, relu, bias, x0

Same as Eq. 19, 20, 21 and with the additional constraint on **W** to respect Dale’s law which constrains hidden units to either act purely excitatory or inhibitory. Here we set 80% of the hidden units to be excitatory, 20% to be inhibitory (implementation inspired by [43]):

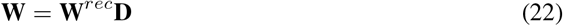

with *W^rec^*(*i, j*) = *max*(0, *W*(*i, j*)), with *i* = 1 : *N, j* = 1 : *N* and a diagonal matrix 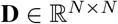 defined as:

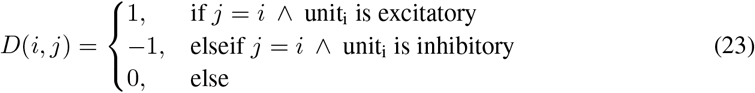

To ensure convergence during training, in this last RNN type **W** was initialized with a Gamma distribution 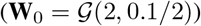.

We trained these alternative RNN types on the context-dependent integrator task. We find that also in these RNNs the inferred operative dimensions successfully identify a low-dimensional functional subspace of the connectivity that is sufficient to solve the task (Fig. 15).

**Figure 15:**
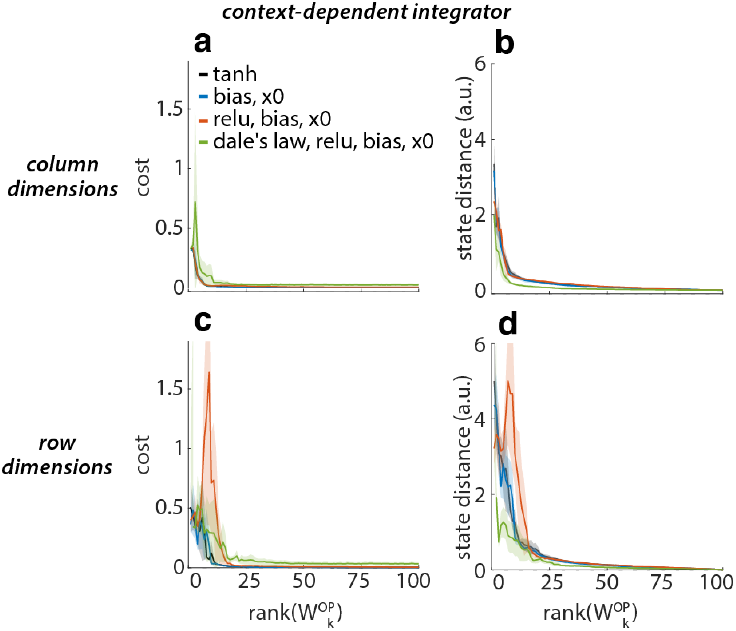
Operative dimensions for various network types. (**a**) Network output cost of networks with reduced-rank weight matrix 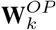 for *k* = 1 : *N* (Eq. 12) for different network types (section A.3.7). (**b**) State distance between trajectories in the full-rank network and in networks with reduced-rank weight matrix 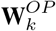 for different network types (section A.3.7). (**c-d**) Same as (**a-b**) for global operative row dimensions. (**a-d**) Averaged over 5 networks per network type; shaded area: *mad* over networks.

#### A.3.8 Operative dimensions applied to sequential MNIST

To validate our definition of operative dimensions to a task more closely related to current AI applications, we estimated operative dimensions on networks trained on sequential MNIST [44]. In this task, the RNN is trained to classify hand-written digits when the individual pixels of each image are provided sequentially over time as an input to the network. For simplicity, the RNN equations and training are analogous to those we employed for the other, simpler tasks considered above (see Eq. 1). Here we increased the size of the RNN’s hidden layer from 100 to 200 units (*N* = 200), while the input is 1-dimensional (*U* = 1). This architecture does not achieve state-of-the-art performance on this task.

The properties of high-variance dimensions *PC*(**W**) for sequential MNIST are similar to those in the simpler tasks above (Fig. 16a-d). The network activity **X** is low-dimensional throughout training while the underlying weight matrix **W** is high-dimensional (Fig. 16a, b; shown for 1 representative example network). Furthermore, sequentially removing *PC*(**W**)_*i*_ from W would imply that the RNN requires more than 175 (out of N=200) dimensions in **W** to achieve the full-rank classification accuracy (Fig. 16a, b; results obtained on test set; full-rank classification accuracy=94% on training and test set).

**Figure 16:**
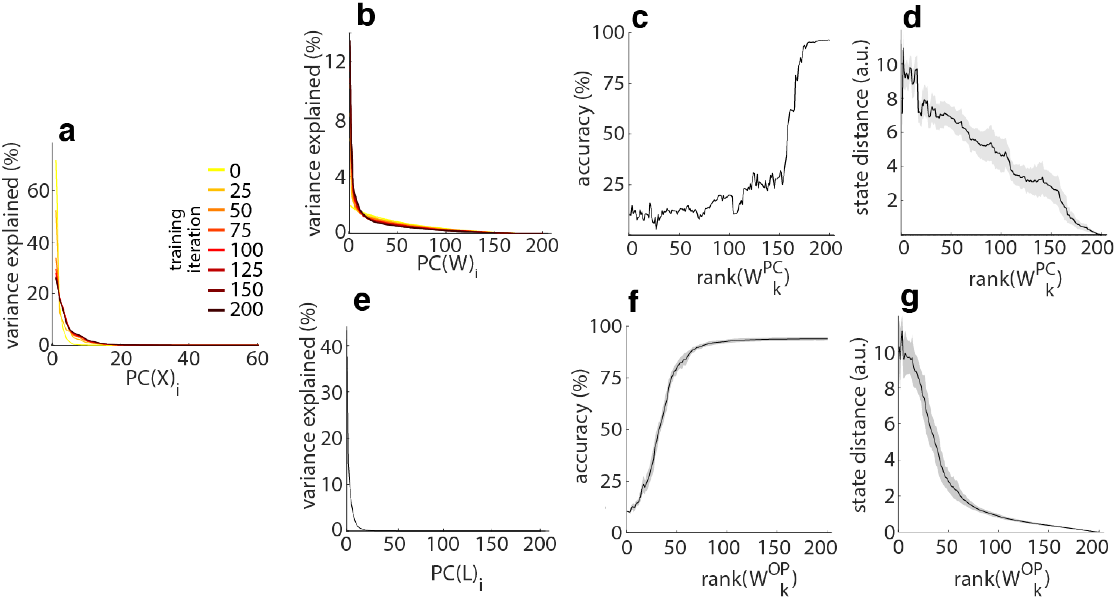
Operative dimensions in sequential MNIST. (**a**) Variance explained (in activity space) by individual PCs of the network activity **X**, shown at different stages of training (test set). (**b**) Variance explained (in weight space) by individual PCs of the weight matrix **W** at different stages of training. (**c**) Network classification accuracy of networks with reduced-rank weight matrices 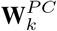 for *k* = 1 : *N* (Eq. 5), (**d**) State distance between trajectories in the full-rank network and in networks with reduced-rank weight matrix 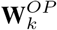. (**e**) Rank of global operative column dimensions, estimated with PC analysis on concatenated local operative dimensions (Eq. 10 and 11). (**f**) Network classification accuracy of networks with reduced-rank weight matrix 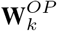 for *k* = 1 : *N* (Eq. 12). (**g**) State distance between trajectories in the full-rank network and in networks with reduced-rank weight matrix 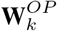. (**a-d**) 1 representative network. (**e-g**) Based on global operative *column* dimensions and averaged over 10 networks per task; shaded area: *mad*.

To identify a functionally relevant subspace in **W** we proceeded as above. We collected the local operative dimensions at *P* = 3950 sampling locations that were equally spaced in time along a random subset of trials (at every 10th time step along 50 randomly selected trials of the training set). Placing sampling locations along the condition average trajectories (averaged over all trials of the same output class) yielded slightly worse performance, most likely due to a lower number of possible sampling locations. Overall, sequential MNIST required a higher number of sampling locations than the simpler tasks presented above [19, 20].

The resulting operative dimensions reveal that the RNN trained on sequential MNIST requires only 58 dimensions (out of 200) to perform the task with the original classification accuracy (here we define *original classification accuracy* as 95% of the accuracy of the corresponding full-rank RNN; Fig. S16; averaged over 10 networks). The operative dimensions thus identify a functionally relevant subspace that is of substantially lower dimensionality than the full-rank weight matrix **W** (results for column dimensions). Further analysis using output-class-specific operative dimensions might reveal valuable insights into the computation implemented by these networks.

#### A.3.9 Local, linear dynamics in reduced-rank RNN using operative dimensions

Computations in RNNs can often be understood by analyzing linear approximations of the dynamics around fixed points or slow points [20]. Here we ask how well the reduced-rank approximations we derived can approximate the local linearized dynamics of the full-rank networks. We find that the operative dimensions of the connectivity are sufficient to reproduce the dominant local, linear dynamics in the full-rank RNNs, reinforcing the finding that the reduced-rank RNNs capture the key computations of the full-rank or closest reduced-rank system.

Here we study the local, linear dynamics in the context-dependent integrator and the sine wave generator by linearizing around slow points of the full-rank RNN 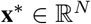 located on the condition average trajectories [20, 19]. We obtain a linear system approximation 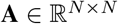:

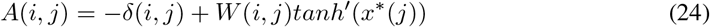

with *i* = 1 : *N, j* = 1 : *N* and

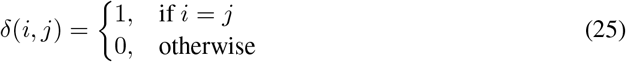

Similarly, we obtain the linear system approximations 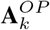 for the reduced-rank RNN using 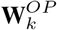 with *k* = 1 : *N* (Eq. 12):

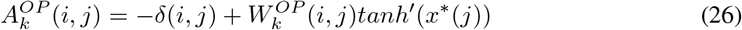

with *i* = 1 : *N, j* = 1 : *N*.

These linear system approximations 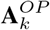 are then studied by analyzing their eigenvalue decomposition:

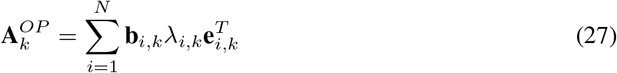

with 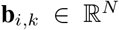 as the i-th right eigenvector, 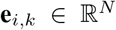 as the i-th left eigenvector and λ_*i,k*_ the i-th eigenvalue of 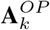 with rank *k*. The full-rank 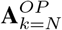 consists of one dominant eigenvector 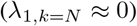 with the remaining modes fast decaying (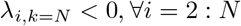, as described in [19]). To compare the eigenvectors and eigenvalues of the reduced-rank systems 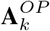 to each other we have to ensure a consistent sorting of their values across linearized systems 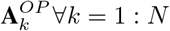. Therefore we sorted the eigenvalues of the full-rank system 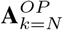 in descending order of the absolute eigenvalues and then used matlab’s *eigenshuffle* (https://www.mathworks.com/matlabcentral/fileexchange/22885-eigenshuffle) to sort the remaining reduced-rank systems to be as similar as possible to the full-rank system.

To measure the similarity between the full-rank and reduced-rank linear dynamics we considered the following quantities (Fig. S17):

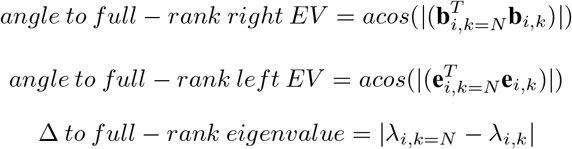

Note that the linearization is performed at the location of slow points **x**\* of the full-rank RNN, which are not necessarily slow points of the dynamics in the reduced-rank RNN. This implies that in the reduced-rank networks the linearized dynamics can be expected to approximate the non-linear dynamics less well than in the full-rank network (or only at some distance from **x**\*, see [20]). Despite this limitation, we find that the inferred dominant linear dynamics is largely preserved in the reduced-rank RNN. In reduced-rank RNN based on only the first few global operative dimensions, the first right eigenvector, left eigenvector and eigenvalue remain very close to their original values in the full-rank RNN; the fast decaying modes instead require the full-rank weight matrix to be retrieved (Fig. S17).

**Figure 17:**
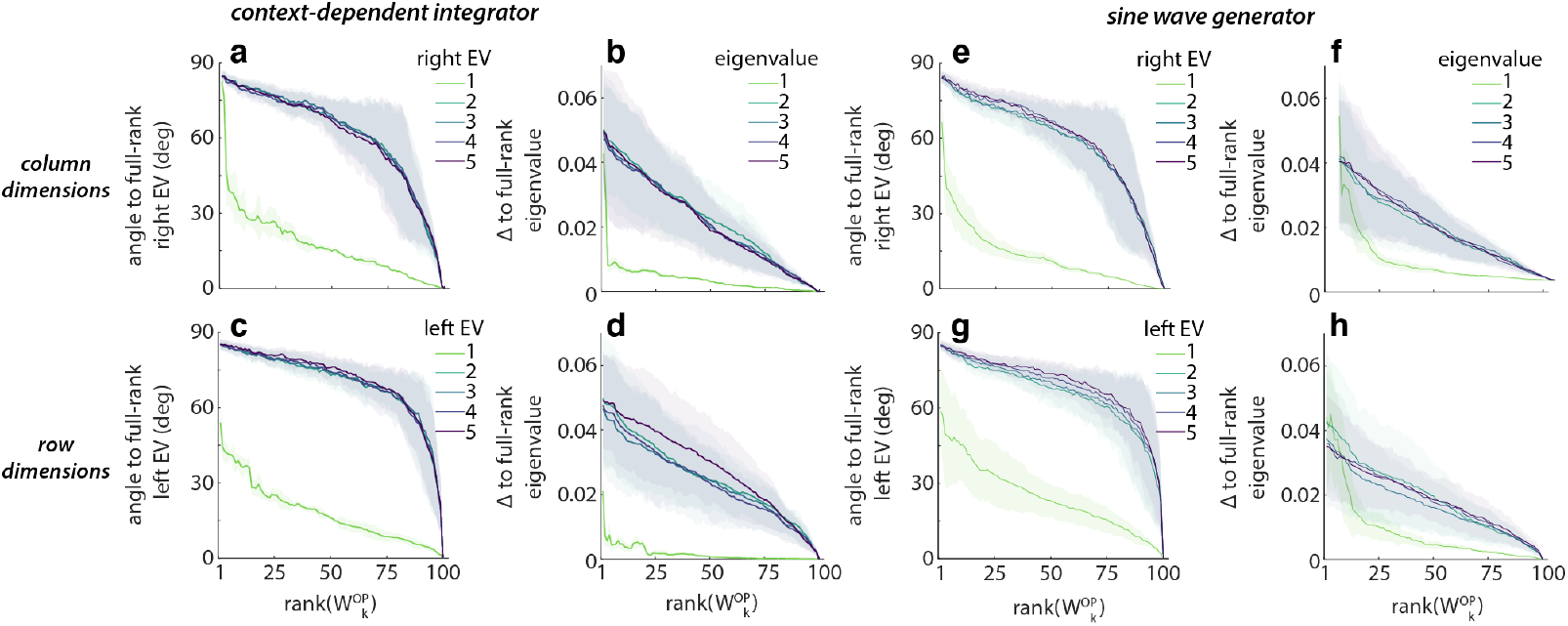
Local, linear dynamics in reduced-rank networks. (**a**) Subspace angle between the right eigenvectors of full-rank RNN and the right eigenvectors of reduced-rank RNN with 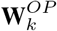 for *k* = 1 : *N*. (**b**) Absolute difference between the eigenvalues of full-rank RNN and the eigenvalues of reduced-rank RNN with 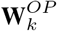 for *k* = 1 : *N*. (**c**) Subspace angle between the left eigenvectors of full-rank RNN and the left eigenvectors of reduced-rank RNN with 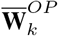 for *k* = 1 : *N* (global operative row dimensions). (**d**) Absolute difference between the eigenvalues of full-rank RNN and the eigenvalues of reduced-rank RNN with 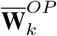 for *k* = 1 : *N* (global operative row dimensions). (**a** - **d**) For context-dependent integration RNN; shaded area: *mad* over *P* = 120 sampling locations of 1 representative network. (**e** - **h**) Same as (**a** - **d**) for sine wave generator with *P* = 121.

#### A.3.10 Operative dimensions for different number of sampling locations

To accurately identify the functionally relevant subspace in **W** it is crucial to define appropriate and sufficient sampling locations **y**_*t*_ to collect the local operative dimensions at. To illustrate how the inferred operative dimensions change depending on the number of sampling locations, we systematically reduced the number of sampling locations used to generate the global operative dimensions while keeping the sampling locations equally distributed over all condition average trajectories (Fig. S18). The global operative dimensions of the context-dependent integration networks are still accurate with fewer sampling locations, whereas the global operative dimensions of the sine wave generator networks generally require more sampling locations to properly capture the functionally relevant subspace in **W**.

**Figure 18:**
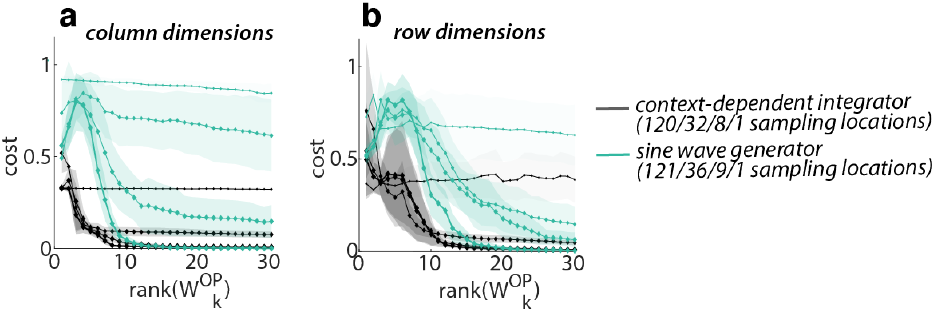
Operative dimensions for different number of sampling locations. (**a**) Network output cost of networks with reduced-rank weight matrix 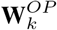 for *k* = 1 : *N* (Eq. 12) for network trained on context-dependent integration using different number of sampling locations. (**b**) Same as (**a**) for global operative *row* dimensions. (**a** - **b**) Averaged over 20 networks per task; shaded area: *mad* over networks; size of diamond-markers corresponds to indicated number of sampling locations (see legend).

#### A.3.11 Single unit contributions to the functional subspace of the connectivity

Our analysis using function-specific operative dimensions (section 2.2.3) showed for the contextdependent integration network that different functional modules are implemented using distinct subspaces in **W**. Functionally relevant weight subspaces are shared between choice_1_ and choice_2_, but not between context1 and context2 (Fig. 5). Here we ask if this functional specificity of operative dimensions is reflected also at the level of the connectivity of single units. Specifically, we focus on two simple properties of a given network unit, namely its effective output and total recurrent input. We define these two quantities for specific task conditions and times by exploiting the inferred operative dimensions.

We consider the effective output and total recurrent input of each unit at every sampling location **y**_*j*_ (*j* = 1 : *P*) separately by generating one reduced-rank approximations 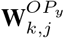 (*k* = 1 : *N*, *j* = 1 : *P*) per sampling location. Every 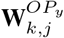 consists of only the local operative dimensions defined for the respective sampling location **y**_*j*_ and thereby provides a reduced-rank approximation that is tailored to perform only specific parts of the network function, similarly to our approach in in defining function-specific operative dimensions (section 2.2.3). Using these 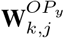 allows us to isolate the input and output of each unit that is functionally relevant to solve specific network functions, i.e. to reproduce activity at specific times and conditions.

We define the total effective output that unit *i* sends to all other hidden units as the norm of the weight matrix column *i* scaled by the network activity of unit *i* at sampling location **y**_*j*_ (*tanh*(*y_t_*(*i*)) = *r_t_*(*i*)):

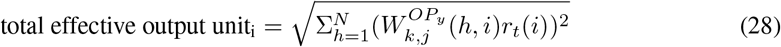

This total effective output is defined separately for each sampling location **y**_*j*_. Here *k* = 1, as the operative column dimensions at every location are always rank 1 (*k* = 1; see section A.2.3).

Similarly, we define the total *recurrent* input received by each unit *i* from all other hidden units as the dot product of the weight matrix row *i* with the network activity **r**_*t*_ = *tanh*(**y**_*t*_).

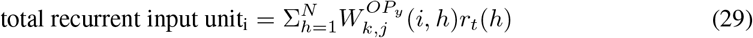

Again, this recurrent input is defined separately for each sampling location **y**_*j*_. Here *k* was set to include only functionally relevant local operative row dimensions for which Δ*f* > 10^−6^ (*k* ≈ 10).

These total recurrent inputs and effective outputs are shown in Fig. S19a, d) for each unit (x-axis) and time/condition (y-axis). This plot does not reveal any obvious structure. Specifically, we find no evidence that particular units are contributing to the functional inputs or outputs preferentially in particular conditions but not others (e.g. context or choice).

In addition to the above unit properties, which combine information about the reduced-rank weight matrix 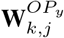 and the network activities **r**_*t*_, we also considered simpler properties based only on 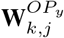, i.e. only on the units’ connectivity. Specifically, we analyzed the norm of each column and each row in the 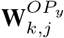, at every sampling location **y**_*t*_, as an alternative measure of the contribution of each unit to specific network functions (Fig. S19b-c, e-f). However, similar to Fig. 19a, d), these measures do not show any obvious structure over time and conditions at the level of single units.

**Figure 19:**
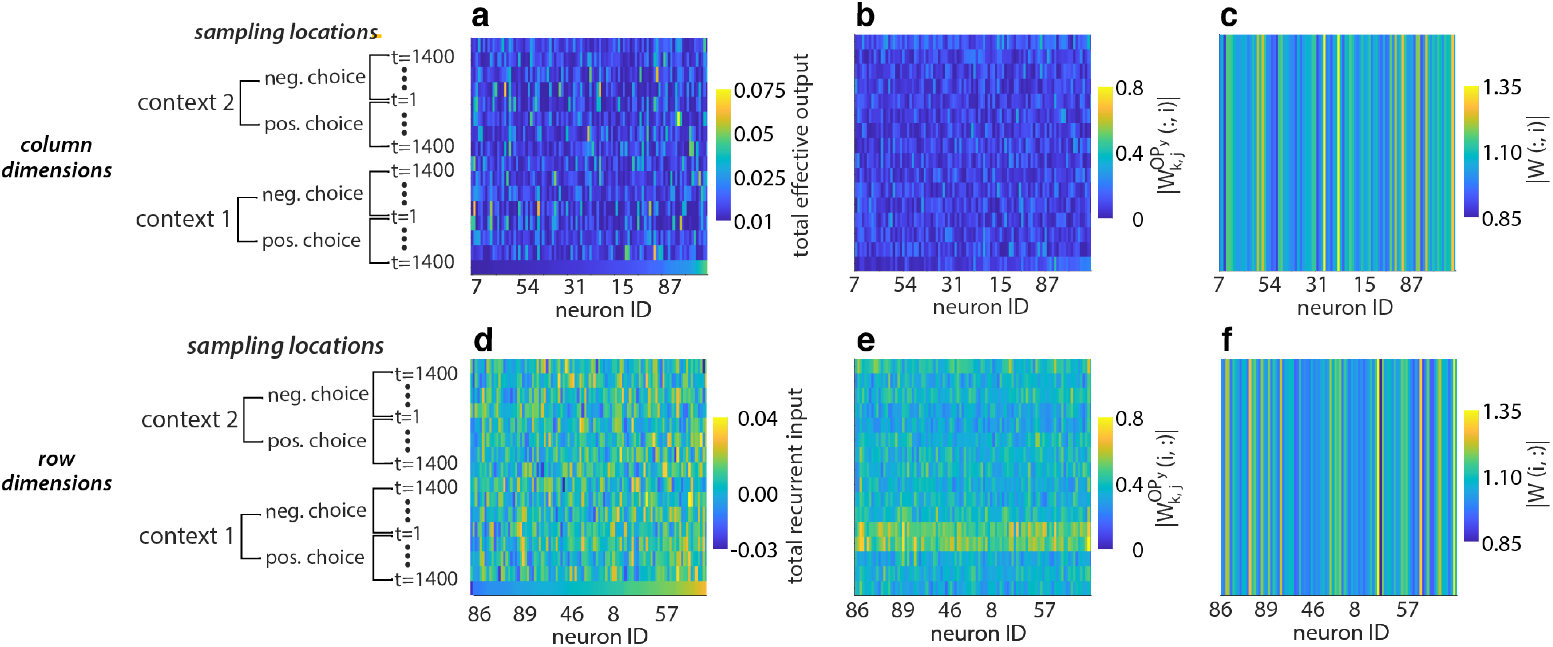
Functional contribution of individual network units. (**a**) Total effective output of each unit *i* at sampling location **y**_*j*_ as defined in Eq. 28. (**b**) Norm of column *i* in reduced-rank weight matrix 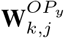 at different sampling locations **y**_*j*_ (i corresponding to neuron ID along x-axis). (**c**) Norm of column *i* in full-rank weight matrix **W** at different sampling locations **y**_*j*_ (i corresponding to neuron ID along x-axis). (**d**) Total recurrent input of each unit *i* at every sampling location **y**_*j*_ as defined in Eq. 28. (**e**) Norm of row *i* in reduced-rank weight matrix 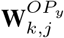 at different sampling locations **y**_*j*_ (i corresponding to neuron ID along x-axis). (**f**) Norm of row *i* in in full-rank weight matrix **W** at different sampling locations **y**_*j*_ (i corresponding to neuron ID along x-axis). (**a-c**) sorted by values of last row in (**a**). (**d-f**) sorted by values of last row in (**d**). (**a-f**) Sampling locations subsampled, shown are locations with congruent inputs at t=1, 400, 900, 1400.

Overall these observations suggest that all neurons are at least partially involved in creating the functionally relevant RNN dynamics at all times and in all conditions. This becomes apparent in Fig. 19a-b, d-e) as the values along every column (corresponding to the total recurrent input or effective output per unit) change abruptly between sampling locations from similar times and conditions (y-axis). However, more detailed analysis of the reduced-rank weight matrices and network activities may well reveal structure at the level of units that is not apparent from the simple properties that we analyzed here.

#### A.3.12 Alignment between global operative column and row dimensions

The global operative column and row dimensions both span a low-dimensional subspace in the weight matrix **W** which is sufficient for the RNNs to perform the task (Fig. 3b-g). However, the two subspaces show little similarity. In the context-dependent integration network, only the first global operative column dimensions are weakly aligned to the first operative row dimensions. The remaining dimensions show no alignment to each other (Fig. S20a). Similarly in the sine wave generator networks, the global operative column and row dimensions show little similarity (Fig. S20b).

Overall, operative column and row dimensions provide complementary insights into RNN computations. In broad terms, the column dimensions in **W** describe the *output connections* of each hidden unit, whereas the row dimensions describe the *input connections* of each hidden unit. The respective operative dimensions in turn identify the functionally relevant subspaces in the network connectivity. If these subspaces of the network connectivity are interpreted as subspaces in the network activity, they might provide a tool to compare each unit’s functionally relevant input and output subspaces, i.e. to determine how the activity of a given hidden unit is shaped by, and how it influences, the activity in the remainder of the network.

**Figure 20:**
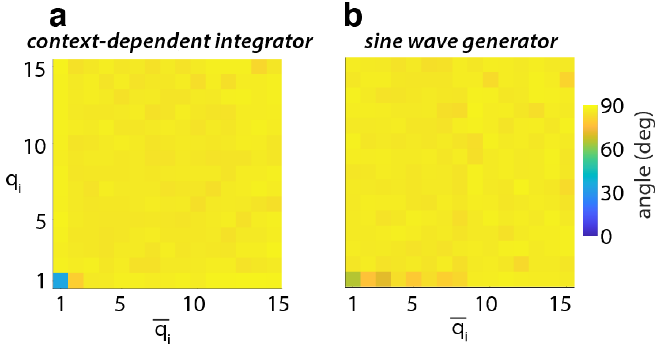
Alignment between global operative column and row dimensions. (**a**) Subspace angle between global operative column dimensions **q**_*i*_ and global operative row dimensions 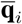 for contextdependent integration networks. (**b**) Same as (**a**) for sine wave generation networks. (**a-b**) Averaged over 20 networks.

#### A.3.13 Alignment of operative dimensions over sampling locations

The global operative dimensions are a combination of the local operative dimensions of all sampling locations. To illustrate the large differences between local operative dimensions across sampling locations, here we show how the local operative dimensions from different sampling locations are aligned to each other. We find that local operative dimensions tend to be similar if their sampling locations are close to each other in state space. However, more distant sampling locations generally yield almost orthogonal local operative dimensions (Fig. S21b, e). Similarly, we test the alignment between the local operative dimensions at a particular sampling location and the global operative dimensions. We find that the global operative dimensions are not preferentially aligned to any local operative dimensions defined at particular sampling location, but rather are partially aligned to the local operative dimensions from all sampling locations (Fig. S21c, f, i, l).

**Figure 21:**
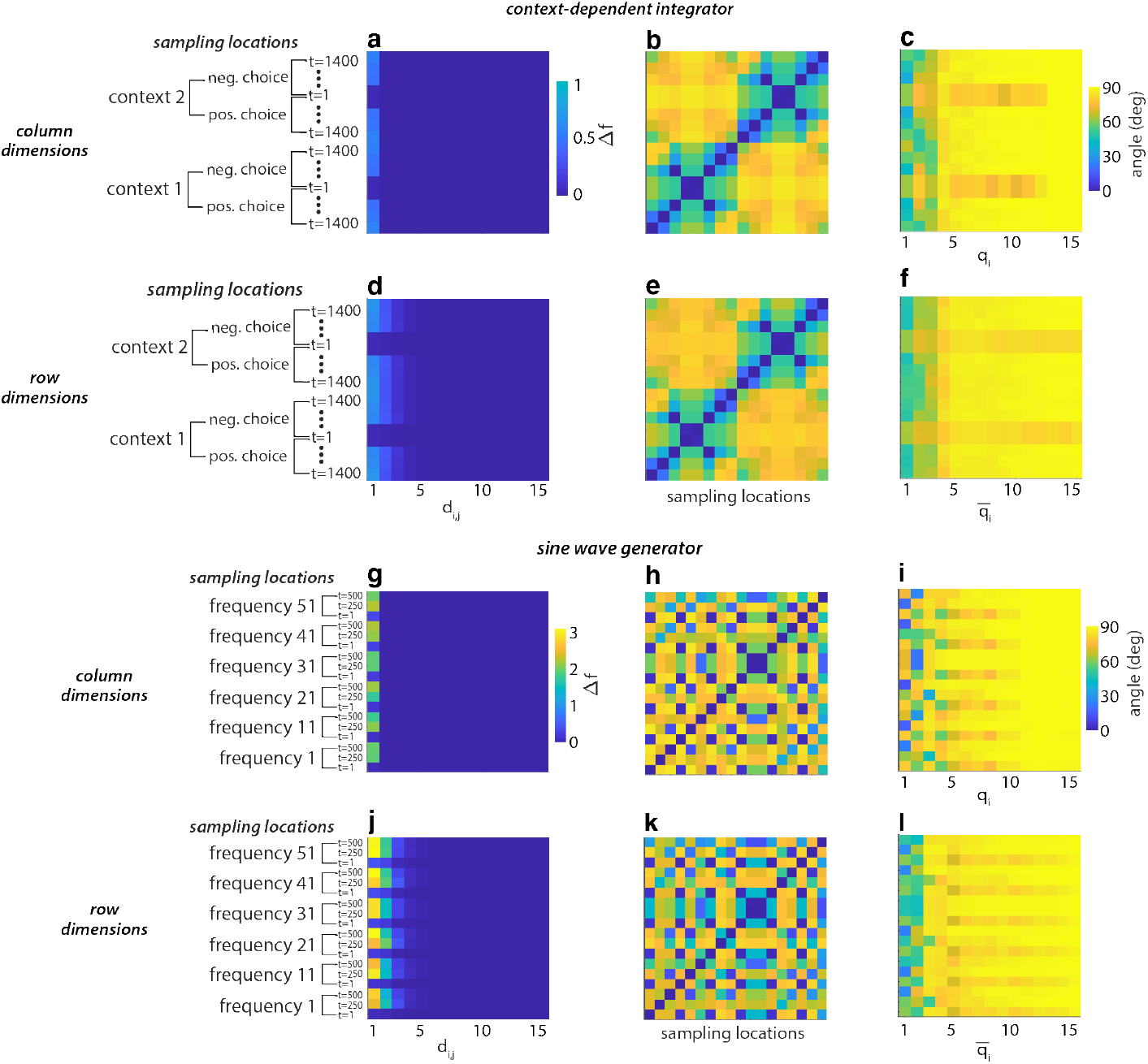
Alignment of operative dimensions over sampling locations. (**a**) Δ*f* for local operative column dimensions of context-dependent integrator. The subspace of local operative column dimensions is 1-dimensional at all sampling locations. (**b**) Pairwise subspace angle of first local operative column dimensions across sampling locations. Local operative column dimensions gradually change over state space, with closer sampling locations yielding more similar local operative column dimensions. (**c**) Subspace angle between the first local operative column dimension at each sampling location and the global operative column dimensions. The first few global operative column dimensions are partially aligned with local operative column dimensions from most sampling locations. (**d-f**) Same as (**a-c**) for operative *row* dimensions for context-dependent integrators. (**g-i**) Same as (**a-c**) for operative *column* dimensions for sine wave generators. (**j-l**) Same as (**a-c**) for operative *row* dimensions for sine wave generators. (**a-f**) Sampling locations are sorted based on the spatial proximity to each other, moving along the line attractor over time in each context. (**g-l**) Sampling locations are sorted based on the input frequency and then the time along the respective condition average trajectory, showing that local operative dimensions are not shared per frequency. (**a-l**) Averaged over 20 networks; only a subsample of all sampling locations shown; all subspace angles are computed considering only the first local operative column or row dimensions at each sampling location.

## References

[1] Peiran Gao and Surya Ganguli. On simplicity and complexity in the brave new world of large-scale neuroscience. Current opinion in neurobiology, 32:148–155, 2015.

[2] Danielle S Bassett and Olaf Sporns. Network neuroscience. Nature neuroscience, 20(3): 353–364, 2017.

[3] Omri Barak. Recurrent neural networks as versatile tools of neuroscience research. Current opinion in neurobiology, 46:1–6, 2017.

[4] David Sussillo. Neural circuits as computational dynamical systems. Current Opinion in Neurobiology, 25:156–163, 2014. ISSN 0959-4388. doi: https://doi.org/10.1016/j.conb.2014.01.008. URL https://www.sciencedirect.com/science/article/pii/S0959438814000166. Theoretical and computational neuroscience.

[5] L.F. Abbott. Theoretical neuroscience rising. Neuron, 60(3):489–495, 2008. ISSN 0896-6273. doi: https://doi.org/10.1016/j.neuron.2008.10.019. URL https://www.sciencedirect.com/science/article/pii/S0896627308008921.

[6] John J Hopfield. Neural networks and physical systems with emergent collective computational abilities. Proceedings of the national academy of sciences, 79(8):2554–2558, 1982.

[7] Rani Ben-Yishai, R Lev Bar-Or, and Haim Sompolinsky. Theory of orientation tuning in visual cortex. Proceedings of the National Academy of Sciences, 92(9):3844–3848, 1995.

[8] Elizabeth Gardner. The space of interactions in neural network models. Journal of physics A: Mathematical and general, 21(1):257, 1988.

[9] Carl Van Vreeswijk and Haim Sompolinsky. Chaos in neuronal networks with balanced excitatory and inhibitory activity. Science, 274(5293):1724–1726, 1996.

[10] Francesca Mastrogiuseppe and Srdjan Ostojic. Intrinsically-generated fluctuating activity in excitatory-inhibitory networks. PLoS computational biology, 13(4):e1005498, 2017.

[11] Johnatan Aljadeff, Merav Stern, and Tatyana Sharpee. Transition to chaos in random networks with cell-type-specific connectivity. Physical review letters, 114(8):088101, 2015.

[12] Haim Sompolinsky, Andrea Crisanti, and Hans-Jurgen Sommers. Chaos in random neural networks. Physical review letters, 61(3):259, 1988.

[13] Alexander Rivkind and Omri Barak. Local dynamics in trained recurrent neural networks. Physical review letters, 118(25):258101, 2017.

[14] Andrew M Saxe, James L McClelland, and Surya Ganguli. Exact solutions to the nonlinear dynamics of learning in deep linear neural networks. arXiv preprint arXiv:1312.6120, 2013.

[15] Nadav Timor, Gal Vardi, and Ohad Shamir. Implicit regularization towards rank minimization in relu networks. arXiv preprint arXiv:2201.12760, 2022.

[16] Carina Curto, Jesse Geneson, and Katherine Morrison. Fixed points of competitive threshold-linear networks. Neural computation, 31(1):94–155, 2019.

[17] Francesca Mastrogiuseppe and Srdjan Ostojic. Linking connectivity, dynamics, and computations in low-rank recurrent neural networks. Neuron, 99(3):609–623, 2018.

[18] Alexis Dubreuil, Adrian Valente, Manuel Beiran, Francesca Mastrogiuseppe, and Srdjan Ostojic. Complementary roles of dimensionality and population structure in neural computations. biorxiv, 2020.

[19] Valerio Mante, David Sussillo, Krishna V Shenoy, and William T Newsome. Context-dependent computation by recurrent dynamics in prefrontal cortex. nature, 503(7474):78–84, 2013.

[20] David Sussillo and Omri Barak. Opening the black box: low-dimensional dynamics in highdimensional recurrent neural networks. Neural computation, 25(3):626–649, 2013.

[21] Niru Maheswaranathan, Alex Williams, Matthew Golub, Surya Ganguli, and David Sussillo. Universality and individuality in neural dynamics across large populations of recurrent networks. Advances in neural information processing systems, 32, 2019.

[22] Friedrich Schuessler, Francesca Mastrogiuseppe, Alexis Dubreuil, Srdjan Ostojic, and Omri Barak. The interplay between randomness and structure during learning in rnns. Advances in neural information processing systems, 33:13352–13362, 2020.

[23] Peiran Gao, Eric Trautmann, Byron Yu, Gopal Santhanam, Stephen Ryu, Krishna Shenoy, and Surya Ganguli. A theory of multineuronal dimensionality, dynamics and measurement. BioRxiv, page 214262, 2017.

[24] Jimmy Smith, Scott Linderman, and David Sussillo. Reverse engineering recurrent neural networks with jacobian switching linear dynamical systems. Advances in Neural Information Processing Systems, 34:16700–16713, 2021.

[25] Bart Besselink, Umut Tabak, Agnieszka Lutowska, Nathan Van de Wouw, H Nijmeijer, Daniel J Rixen, ME Hochstenbach, and WHA Schilders. A comparison of model reduction techniques from structural dynamics, numerical mathematics and systems and control. Journal of Sound and Vibration, 332(19):4403–4422, 2013.

[26] Peter Benner, Serkan Gugercin, and Karen Willcox. A survey of projection-based model reduction methods for parametric dynamical systems. SIAM review, 57(4):483–531, 2015.

[27] Ivan Ivanov, Ranadip Pal, and Edward R Dougherty. Dynamics preserving size reduction mappings for probabilistic boolean networks. IEEE Transactions on Signal Processing, 55(5): 2310–2322, 2007.

[28] Aurélien Naldi, Elisabeth Remy, Denis Thieffry, and Claudine Chaouiya. Dynamically consistent reduction of logical regulatory graphs. Theoretical Computer Science, 412(21):2207–2218, 2011.

[29] Francesco Ginelli, Pietro Poggi, Alessio Turchi, Hugues Chaté, Roberto Livi, and Antonio Politi. Characterizing dynamics with covariant lyapunov vectors. Physical review letters, 99 (13):130601, 2007.

[30] Christopher L Wolfe and Roger M Samelson. An efficient method for recovering lyapunov vectors from singular vectors. Tellus A: Dynamic Meteorology and Oceanography, 59(3): 355–366, 2007.

[31] Zoltan Toth and Eugenia Kalnay. Ensemble forecasting at nmc: The generation of perturbations. Bulletin of the american meteorological society, 74(12):2317–2330, 1993.

[32] Niru Maheswaranathan, Alex Williams, Matthew Golub, Surya Ganguli, and David Sussillo. Reverse engineering recurrent networks for sentiment classification reveals line attractor dynamics. Advances in neural information processing systems, 32, 2019.

[33] Feng-Lei Fan, Jinjun Xiong, Mengzhou Li, and Ge Wang. On interpretability of artificial neural networks: A survey. IEEE Transactions on Radiation and Plasma Medical Sciences, 5(6): 741–760, 2021.

[34] Grégoire Montavon, Wojciech Samek, and Klaus-Robert Müller. Methods for interpreting and understanding deep neural networks. Digital Signal Processing, 73:1–15, 2018.

[35] Quan-shi Zhang and Song-Chun Zhu. Visual interpretability for deep learning: a survey. Frontiers of Information Technology & Electronic Engineering, 19(1):27–39, 2018.

[36] Lea Duncker, Laura Driscoll, Krishna V Shenoy, Maneesh Sahani, and David Sussillo. Organizing recurrent network dynamics by task-computation to enable continual learning. Advances in neural information processing systems, 33:14387–14397, 2020.

[37] Genta Indra Winata, Andrea Madotto, Jamin Shin, Elham J Barezi, and Pascale Fung. On the effectiveness of low-rank matrix factorization for lstm model compression. arXiv preprint arXiv:1908.09982, 2019.

[38] James Martens and Ilya Sutskever. Learning recurrent neural networks with hessian-free optimization. In ICML, 2011.

[39] Sepp Hochreiter and Jürgen Schmidhuber. Long short-term memory. Neural computation, 9(8): 1735–1780, 1997.

[40] Kyunghyun Cho, Bart Van Merriënboer, Dzmitry Bahdanau, and Yoshua Bengio. On the properties of neural machine translation: Encoder-decoder approaches. arXiv preprint arXiv:1409.1259, 2014.

[41] Herbert Jaeger. Adaptive nonlinear system identification with echo state networks. Advances in neural information processing systems, 15, 2002.

[42] Wolfgang Maass, Thomas Natschläger, and Henry Markram. Real-time computing without stable states: A new framework for neural computation based on perturbations. Neural computation, 14(11):2531–2560, 2002.

[43] H Francis Song, Guangyu R Yang, and Xiao-Jing Wang. Training excitatory-inhibitory recurrent neural networks for cognitive tasks: a simple and flexible framework. PLoS computational biology, 12(2):e1004792, 2016.

[44] Li Deng. The mnist database of handwritten digit images for machine learning research. IEEE Signal Processing Magazine, 29(6):141–142, 2012.

